# Gene loss propensity for metallocarboxypeptidase E in insects is shaped by structural versatility and broader expression of metallocarboxypeptidase D but not functional importance

**DOI:** 10.64898/2026.08.05.742955

**Authors:** Christian Wegener, Julian C. Heitkamp, Vera S. Hunnekuhl

## Abstract

Gene loss is a widespread phenomenon that shapes genome evolution, yet the factors determining why certain genes are repeatedly lost while other functionally related genes are retained remain poorly understood. We addressed this question using the peptide-processing metallocarboxypeptidases carboxypeptidase E (CPE) and carboxypeptidase D (CPD), conserved paralogues that are essential for neuropeptide maturation but strikingly differ in their evolutionary fate: the c*pe* gene has been independently lost in two major insect lineages, whereas *cpd/svr* has been universally retained. Combining gene phylogenetic analyses and functional genetics in the beetle *Tribolium castaneum*, and cross-species rescue experiments in the fly *Drosophila melanogaster*, we show that CPE and CPD retained partially interchangeable enzymatic functions despite considerable differences in structure, organismal importance and expression. Contrary to expectations, *cpe* proved more critical than *cpd/svr* for survival and developmental robustness in *Tribolium*, while simultaneous RNAi-mediated downregulation of both genes caused complete larval lethality, demonstrating only partial functional redundancy. Moreover, beetle CPE partially rescued the lethal loss of *Drosophila* CPD, establishing conserved molecular interchangeability across ∼300 million years of insect evolution. Gene phylogenetic analyses further indicate that bilaterian CPE originated through duplication of the second catalytic domain of an ancestral CPD. Together, our results demonstrate that repeated loss of insect *cpe* cannot be explained by reduced functional importance. Instead, we propose that the structural versatility, broader tissue distribution and multifunctionality of CPD, including its multidomain architecture and splice isoforms, enabled compensation for CPE after gene loss, thereby shaping long-term patterns of gene retention and loss during insect evolution.

## Introduction

The evolution of animal genomes is shaped by expansion of gene families as well as by differential loss of non-essential genes on different phylogenetic levels (Albalat and Cañestro 2016; Fernández and Gabaldón 2020). For the loss of a coding gene sequence to become fixed in a lineage, the gene function must have become dispensable. Main drivers underlying gene loss are adaptive evolution or environmental change that lead to a relaxed selection pressure, or the presence of related genes (paralogues) that can substitute the function (Krylov et al. 2003; Albalat and Cañestro 2016; Sharma et al. 2018). As a result, the functional importance of the gene is decreased and the propensity for gene loss is increased. While the general mechanisms underlying gene loss are increasingly understood due to ever-increasing genomic information, there is a lack of studies that experimentally test the interchangeability between functionally related genes acting in essential pathways (Cadigan et al. 1994; Laugier et al. 2005; Loker and Mann 2022). Moreover for most genes we still do not know why they became lost during evolution.

Neuropeptide signalling is an evolutionary ancient and essential cell-to-cell communication system in animals (Grimmelikhuijzen and Hauser 2012; Jekely 2013; Mirabeau and Joly 2013; Achim and Arendt 2014; Elphick et al. 2018). Neuropeptides are short, genetically encoded signalling molecules that are secreted as neuromodulators or hormones from peptidergic neurons or endocrine cells and coordinate numerous essential biological processes. Notably in insects, the most species-rich and diverse clade among animals (Misof et al. 2014), neuropeptides are involved in coordinating essential processes such as metabolism, development and behaviour (Schoofs et al. 2016; Nässel and Zandawala 2019; Nässel 2024).

In all animals, neuropeptides are synthesized from larger precursor proteins by a set of specific enzymes (Rholam and Fahy 2009; Fricker 2012; Pauls et al. 2014). These include prohormone convertases (PCs (Hoshino and Lindberg 2012)) that cleave the precursors at specific mono- or dibasic sites, and specific metallocarboxypeptidases (carboxypeptidase E (CPE) and carboxypeptidase D (CPD) (Fricker 2025a; Fricker 2025b) that remove the C-terminal mono- or dibasic overhangs resulting from PC cleavage. Mice with a CPE loss-of-function mutation (C*pe^fat/^*^fat^) or a knockout of CPE suffer from deleterious consequences for health and survival (Naggert et al. 1995; Woronowicz et al. 2008; Rodriguiz et al. 2013; Sapio and Fricker 2014; Cawley et al. 2016; Ji et al. 2017). Moreover, *Cpe^fat/fat^* mice show strongly reduced neuropeptide levels and produce unusual C-terminally extended immature neuropeptides (Fricker et al. 1996; Rovere et al. 1996; Che et al. 2001; Che et al. 2005). Still, *Cpe^fat/fat^* mice are able to produce at least some mature neuropeptides at low level, a feature commonly attributed to partial compensation by CPD (Song and Fricker 1995). Unlike CPE, which is mostly expressed in nervous and endocrine tissues (Fricker 2025b), CPD is broadly expressed throughout the body and is involved in the processing of numerous proteins that are shuttled through the secretory pathway (Fricker 2025a).

Homologous genes encoding prohormone convertases and peptide-processing carboxypeptidases are found across the animal kingdom (Husson, Mertens, et al. 2007; Tessmar-Raible 2007; Hammond et al. 2019; Pauls et al. 2019; Fritzsche and Hunnekuhl 2021) and their essential function in the neurosecretory pathway is well conserved between vertebrate and invertebrate models (Sidyelyeva and Fricker 2002; Fricker 2007; Wegener et al. 2011; Hoshino and Lindberg 2012; Pauls et al. 2019). In particular, comparative genomic analyses revealed that the prohormone convertases PC1/3 and PC2 and the carboxypeptidases CPE and CPD are highly conserved in insects. Interestingly, however, the vinegar fly *Drosophila melanogaster* – like other dipterans-has lost two of these enzymes, PC1/3 and CPE (Tessmar-Raible 2007; Pauls et al. 2019; Fritzsche and Hunnekuhl 2021) and relies on a single prohormone convertase dPC2/Amontillado (Siekhaus and Fuller 1999) and a single carboxypeptidase dCPD encoded by the gene *silver (svr*) (Settle et al. 1995) for peptide processing. Mutations in the encoding genes are developmentally lethal and cause severe developmental and physiological deficiencies (Settle et al. 1995; Sidyelyeva and Fricker 2002; Rayburn et al. 2003; Rayburn et al. 2009; Rhea et al. 2010) including strongly impaired neuropeptide processing (Rhea et al. 2010; Wegener et al. 2011; Pauls et al. 2019).

While the general conservation of two prohormone convertases and two peptide-processing carboxypeptidases across insects implies that there is a functional importance for all of them, functional evidence for this hypothesis is lacking. Moreover, it is unclear why *Drosophila* has lost PC1/3 and in particular CPE, the carboxypeptidase most relevant for neuropeptide processing in mammals (Fricker et al. 1996; Rovere et al. 1996; Che et al. 2001; Zhang et al. 2008).

Recently, in the red flour beetle *Tribolium castaneum*, we were able to show that prohormone convertase PC1/3, which is lost in *Drosophila*, plays an essential role in larval growth and is not functionally redundant with its paralogue PC2 (Fritzsche and Hunnekuhl 2021). Moreover, both carboxypeptidases CPD and CPE are conserved in *Tribolium.* Gene predictions show that *Tc-cpd/svr* encodes a large three domain protein (tCPD) similar to vertebrate and other insect CPDs, which occur in different splice variants. *Tribolium* CPE (tCPE) by contrast is a shorter protein with only a single peptidase domain, similar to mammalian and other insect CPEs. The simpler domain architecture and the fact that CPE was independently lost in dipteran and hymenopteran insects (Pauls et al. 2019) suggest that CPE may be functionally redundant with CPD in beetles and other insects that have retained both CPs.

Here, we used the genetically amenable *Tribolium* model to test for functional redundancy between the CP paralogues, and to find answers as to why during evolution CPE but not CPD has been lost in certain insect orders, We first characterised the expression and developmental function of both CPs in *Tribolium.* We then performed a cross-species substitution experiment using the *Tribolium cpe*-sequence to rescue the lethal *Drosophila* dCPD *svr*-mutants. Surprisingly our results for the beetle show that *Tc*-*cpe*, the gene independently lost in major insect lineages (Pauls et al. 2019) holds much greater functional importance than *Tc-*c*pd/svr,* which is not essential for *Tribolium* development or viability. Furthermore, *Tc-cpe* is able to partially substitute the loss of *svr* in *Drosophila*, showing that the molecular function is indeed redundant. Our findings imply that the repeated lineage specific loss of *cpe* and not *cpd* was not facilitated by a lack of functional importance of CPE on an organismic level. Instead, the results suggest that the complex structure of *cpd/svr* with two active CP domains and the resulting ability to produce different splice variants with broader tissue distribution and functions favoured and sustained the repeated loss of the *cpe*-gene during insect evolution.

## Results

### 1 Variants and evolutionary history of CPD and CPE proteins

The *Tribolium* genome contains both a gene for tCPD and tCPE (Pauls et al. 2019). A more thorough analysis now identified two isoforms of the beetle *cpd/svr* gene, which encode a short protein with only one active CP domain, and a long version containing three CP domains (figure 1A). The long version also contains a transmembrane domain followed by a cytosolic tail (predictions based on InterPro (Blum et al. 2025)), suggesting that it is membrane-tethered. Both the short version of *Tribolium* CPD and the CPE protein lack a predicted transmembrane domain. In *Drosophila*, based on gene models resulting from RNA sequence evidence (FlyBase 2025/4 and (Sidyelyeva and Fricker 2002); see table S2 and figure S2) there are five major protein variants that result from alternative splicing of the first exon and the presence of long- and short versions of two variants (figure 1B, table S2). The 1C-long version only has a partial first carboxypeptidase domain and is also lacking the N-terminal signal peptide important for the trafficking of the polypeptide to the secretory pathway (Reznik and Fricker 2001; Sidyelyeva and Fricker 2002). As for the short and long protein versions of *Tribolium*, only the long fly variants carry a transmembrane domain. Additionally, in *Drosophila*, there are two different C-terminal tails, further increasing the number of protein variants to a total of seven (figure 1B). The presence of seven variants is also backed up by the annotation of 11 differentially spliced mRNAs assigned to the *Drosophila svr* locus in Flybase (flybase.org). Hence *Drosophila*, which lacks a *cpe* gene, produces more variants of CPD proteins than *Tribolium*.

**Figure 1:**
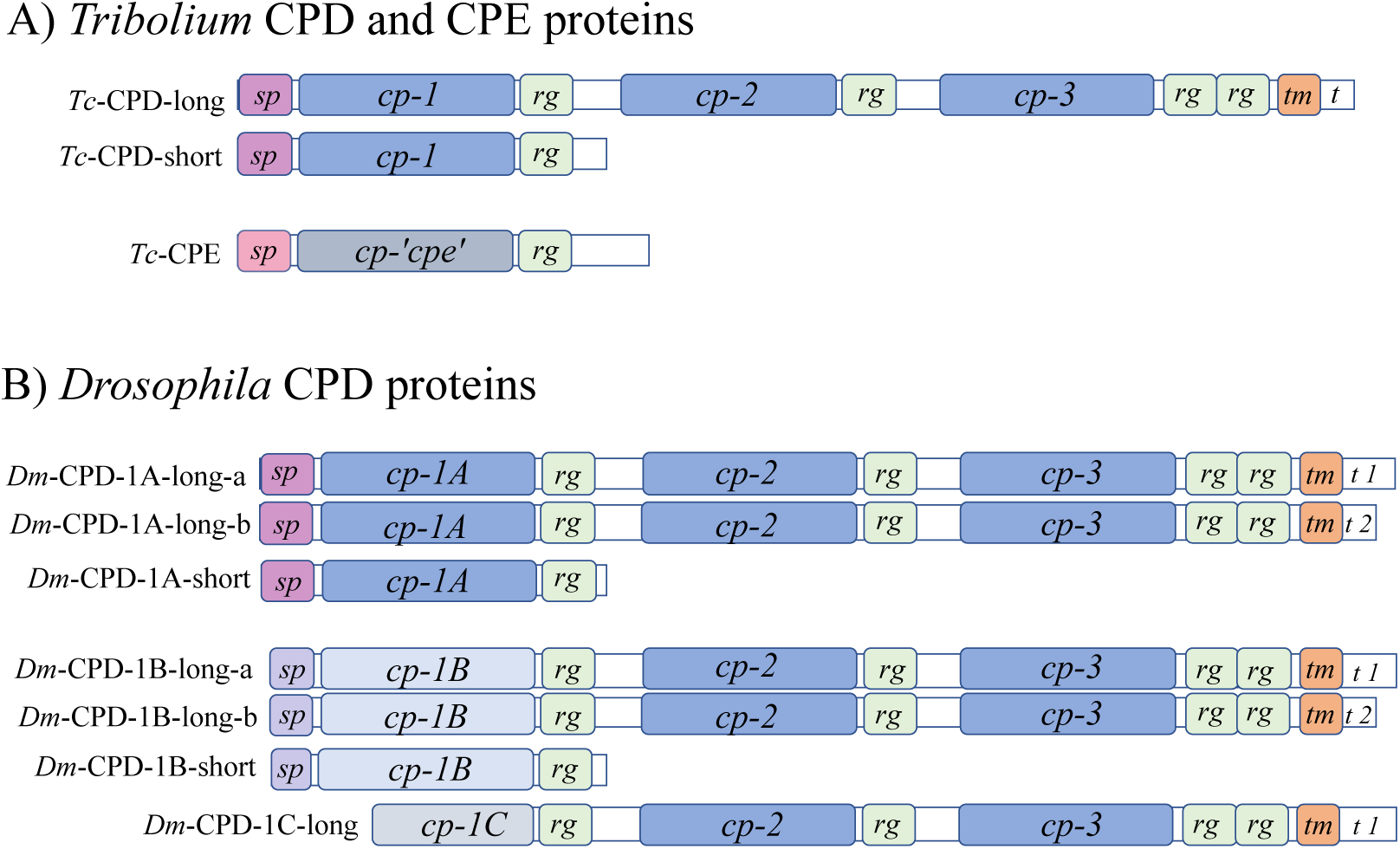
A) Predicted structure of proteins encoded by the short and long *Tc-cpd/svr* isoforms (PCR-validated, see supplementary figure S1 A, B), and single *Tc-cpe* isoform. B) Protein structure of dCPD variant (from flybase.org). sp=signal peptide, cp=carboxypeptidase domain, rg=regulatory domain, tm=transmembrane domain, t=tail.

Interestingly, three-domain *Cpd* genes have been characterized in pre-bilaterian species whereas *Cpe* genes have only been found in bilaterians (Hammond et al. 2019). Both CPE and CPD proteins are closely related members of the M14B subfamily of metallocarboxypeptidases (Fricker 2025a; Fricker 2025b). We therefore hypothesized that *Cpe* originated from a partial duplication of an ancestral *Cpd* gene in the lineage leading to bilaterian animals. We performed a domain specific phylogenetic analysis of CPD and CPE proteins and found that the carboxypeptidase domain of CPEs is most similar to the CPD carboxypeptidase 2 domain. This provides evidence that the *Cpe* gene indeed originates from *Cpd* (figure 2). Notably, we found that the CP2 domains of CPD are more similar to the single CPE domain than to either the CP1 or CP3 domain of CPD (figure 2A), demonstrating a high degree of domain-specific protein sequence conservation.

**Figure 2:**
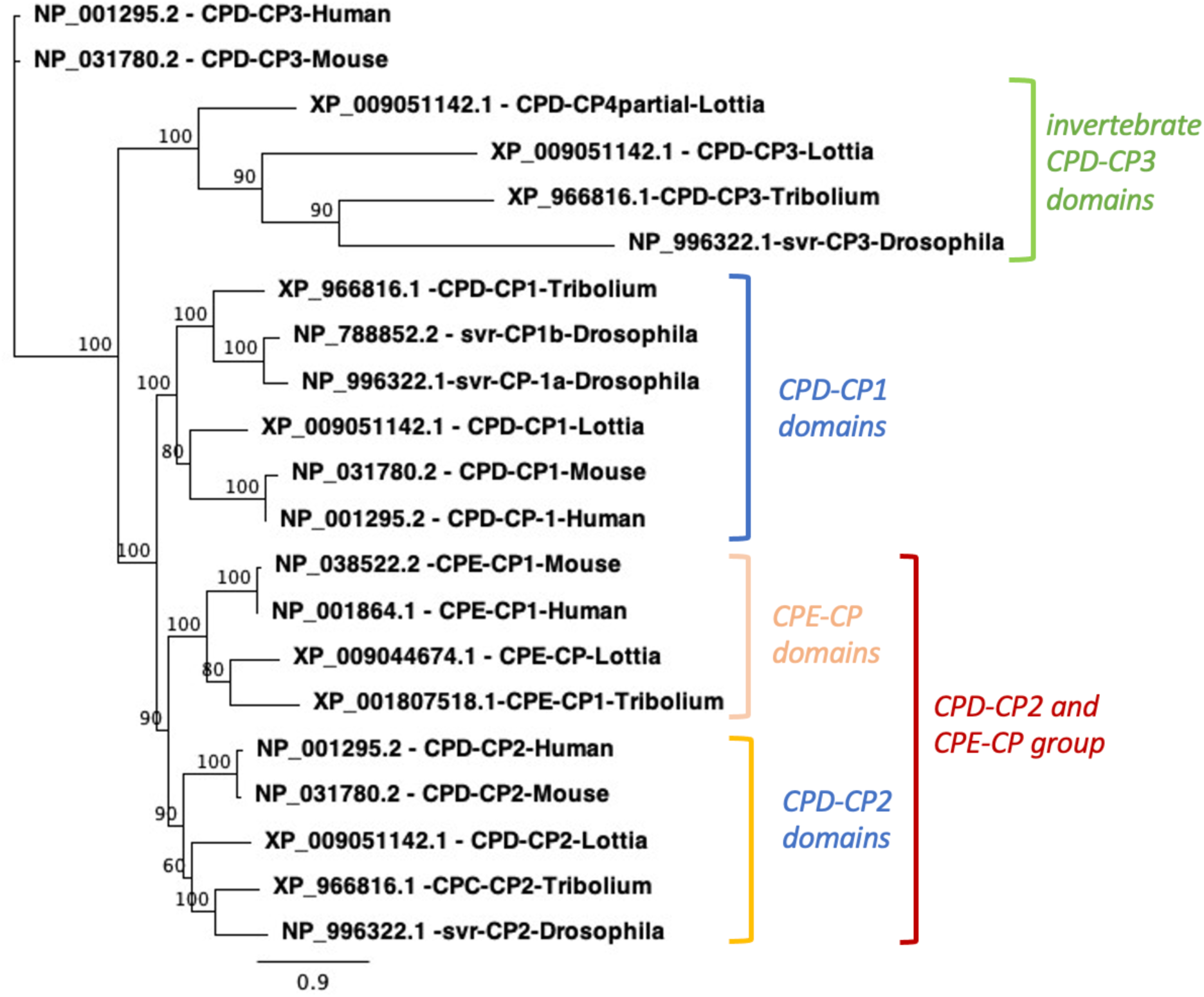
Evolution of *Cpd (svr)* and *Cpe* genes. ML Gene tree (Guindon and Gascuel 2003) based on protein alignment of the individual functional carboxypeptidase domains including sequences from the insects *Tribolium castaneum* and *Drosophila melanogaster*, the mollusc *Lottia gigantea* and the vertebrates mouse and human. Branch labels indicate bootstrap support values (100 repl.). Vertebrate and invertebrate CPE sequences form one group and are more closely related to CPD-CP2 domains than CPD-CP1 or CPD-CP3 domains are. Invertebrate CPD -CP3 domains form a separate branch whereas vertebrate CPD-CP3 domains fail to cluster with one of the groups.

### 2 Only *Tc-cpe* expression shows tissue specificity in the developing and mature nervous system

We examined expression of *Tc-cpd/svr* and *Tc-cpe* over the developmental time course using quantitative PCR on RNA samples extracted from different stages (figure 3 A, B). Expression of *Tc-cpd/svr* was lowest in late embryos (48-72 h) and highest in mobile L7 larvae. However, fold changes ranged only between 0.4 and 1.7, indicating relatively uniform expression levels (figure 3 A). We also performed RNA-*in situ* hybridisation on embryos and larval nervous systems using two different probes for *Tc-cpd/svr*, one that would bind both isoforms and one that would only bind to the long isoform (figure 1, and see suppl. figure S1 C for fragments) but detected no expression. Taken together, these results are suggestive of a uniform expression of *Tc-cpd/svr* below a threshold that is detectable by RNA *in situ* hybridisation. Data on tissue-specific gene expression available on Beetle Atlas (https://motif.mvls.gla.ac.uk/BeetleAtlas/) (Leader et al. 2024) also indicates uniform expression of *Tc-cpd/svr* (acc. # TC015532) across adult and larval tissues.

**Figure 3:**
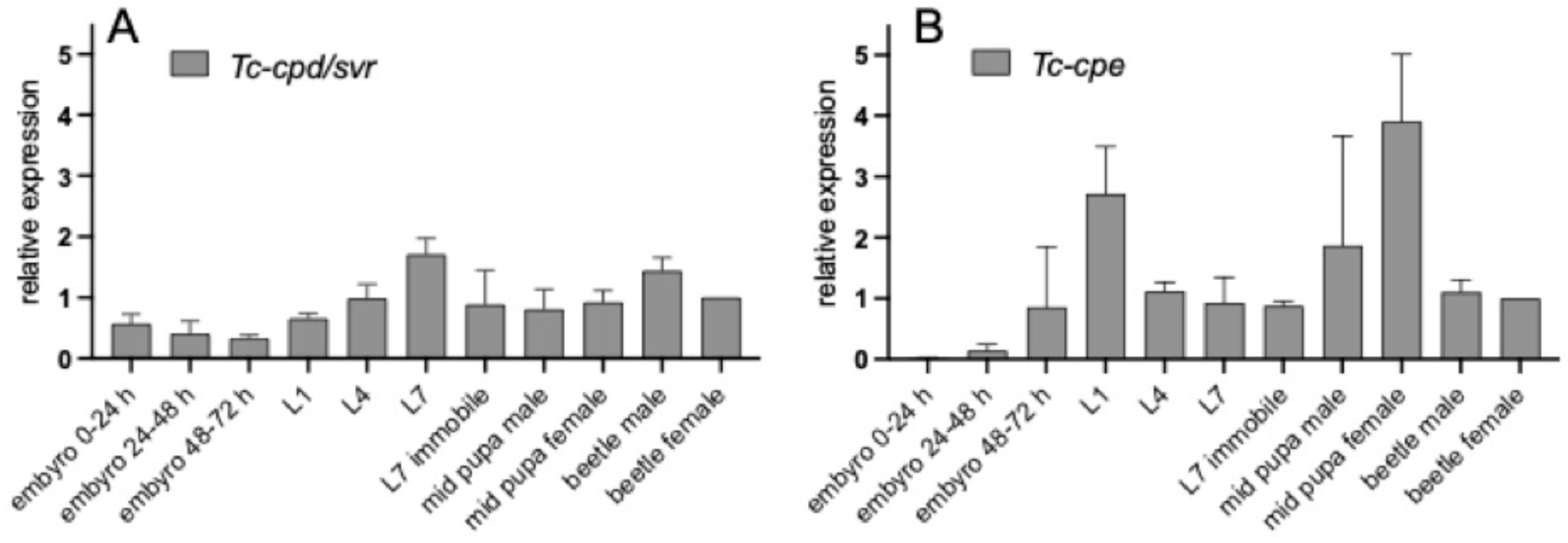
Expression of *Tc-cpd/svr* (A) and *Tc-cpe* (B) across developmental stages. Primers for *Tc-cpd/svr* detect both isoforms (see suppl. figure S1). Data was analysed in relation to the respective ‘female beetle’ samples (≙1) using the ΔΔCT method (Schmittgen and Livak 2008). Error bars show standard deviations (positive values only) across 3 biological repeats.

We next performed qPCR for *Tc-cpe* and found that the gene is not expressed during the earliest embryonic stages (0-24 h). Expression starts from 24-48 h of embryonic development and peaks in L1 larvae and mid-pupae, with highest transcript levels in female pupae (figure 3 B).

The temporal pattern of *Tc-cpe* expression is in line with an expression in the developing central nervous system of the embryo and in the larval nervous system which we confirmed using an RNA-probe against *Tc*-*cpe* (figure 4 A, B). In addition, tissue specific mRNA sequencing data available on Beetle Atlas indicates enrichment of *Tc-cpe (*acc. # TC031795) expression in the larval and adult brain (Leader et al. 2024). In summary, we found evidence that *Tc-cpe* is specifically expressed in the developing and mature nervous system, while *Tc-cpd/svr* expression shows broad expression with little tissue- or temporal specificity.

**Figure 4:**
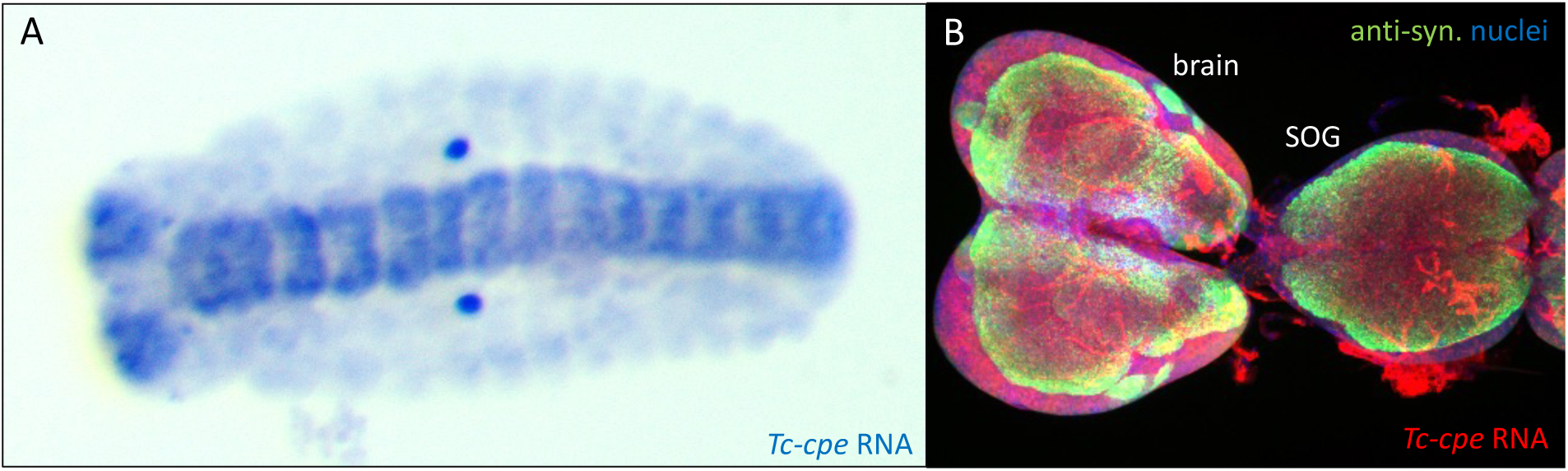
*Tc-cpe* expression detected by RNA-*in situ* hybridisation in an embyro. (A) stage NS 14, expression in the neuroectoderm forming the brain and ventral nervous system (Biffar and Stollewerk 2014) and in the anterior larval nervous system (B). Note that the two strongly stained dots in A are unspecific staining of an embryonic glandular structure and the strongly stained tubular structures in B are trachea which often produce background signal. SOG=suboesophageal ganglion, anti-syn.=anti-synapsin antibody.

### 3 Redundant functions of *Tc-cpe* and *Tc-cpd/svr* during larval development

We performed RNAi experiments to study the functional importance of both *Tc-cpe* and *Tc-cpd/svr* during larval development. There were no sequence matches longer than 8 bp between the used *Tc-cpe* dsRNA and the *Tc-cpd/svr* gene and *vice versa*, excluding possible cross-reactivity between the fragments (with fragments >19 bp considered likely to induce off-target effects (Kulkarni et al. 2006)). We first tested the efficiency of our dsRNAs and found that we achieved very strong knockdown levels of *Tc-cpd/svr* (94 %) and slightly more moderate levels of *Tc-cpe* knockdown (70 %) (figure S3). Previous studies showed that there is no fixed measure as to what level transcripts need to be reduced to interrupt gene function (Mehlhorn et al. 2021) and even though *Tc-cpe* knockdown efficiency was above the 5 % conventional p-value for significance (figure S3), our phenotypic analyses showed, that the knockdown of *Tc-cpe* was sufficient to induce a phenotype. In addition, the low significance level is due to a hight variability of *Tc-cpe*-expression in the control whereas the variability of the knockdown level is in the *Tc-cpe*-RNAi treated specimens is consistently low (figure S3). We found, that in larvae that were injected with *Tc-cpd/svr* or *Tc-cpe* dsRNA at 12 days of age (L4), survival was reduced by 45 % and 54 % respectively (table 1). 23 % of the total *Tc-cpd/svr*-RNAi larvae and 32 % of the *cpe*-RNAi larvae died from an ecdysis phenotype where the larvae were unable to shed their old cuticle. This is a typical phenotype observed in larvae with interrupted neuroendocrine signalling (Rayburn et al. 2003; Fritzsche and Hunnekuhl 2021). To assure a full knockdown of all potential isoforms of *Tc-cpd*, we also included injections of two dsRNA fragments targeting different parts of the gene (figure S1 C). Survival here was only slightly worse than when using only one fragment (table 1). Co-injecting *Tc-cpd/svr*- and *Tc-cpe*-dsRNA at a concentration of 1 µg/µl each (2 µg/µl total) increased the death rate to 100 %, and the larvae all died from the ecdysis phenotype (table 1). To exclude that this was a pure quantitative effect of larger amounts of dsRNA we reduced the concentration to 1 µg/µl (0.5 µg/µl each), equalling the total concentration of dsRNA used in the single knockdowns. We still found a strongly increased occurrence of the lethal ecdysis phenotype (83 %, table 1).

**Table 1:** Mortality and frequency of an ecdysis phenotype in RNAi-treated *Tribolium* L4 larvae.

|  | *dsRed* RNAi<br>1 µg/µl<br>(n= 18) | *Tc-cpd fr 1*<br>µg/µl<br>(n=22) | *Tc-cpd/svr*<br>*fr1 + fr 2*,<br>1 µg/µl<br>(n=12) | *Tc-cpe* 1<br>µg/µl<br>(n=22) | *Tc-cpd/svr +*<br>*cpe*<br>1 µg/µl total<br>(n=12) | *dsRed* RNAi<br>2 µg/µl<br>total (n=12) | *Tc-cpd/svr +*<br>*Tc-cpe*<br>2 µg/µl total<br>(n=12) |
| --- | --- | --- | --- | --- | --- | --- | --- |
| dead | 0 % | 45 % | 50 % | 54 % | 83 % | 8 % | 100 % |
| failed<br>ecdysis | 0 % | 23 % | 25 % | 32 % | 83 % | 0 % | 100 % |

Looking at growth in *Tc-cpe* and *Tc-cpd/svr* RNAi-treated larvae and control groups (figure 5) we found that larvae treated with only *Tc*-*cpd*-dsRNA showed a very slight reduction of growth. The effect on growth was stronger in *Tc-cpe*-dsRNA-treated larvae with individual specimens that failed to reach the required weight for pupation and hence remained in the larval phase for a prolonged period before they died as larvae. Levels of growth reduction in the two-fragment knockdown of *Tc-cpd/svr* were stronger than when using only one fragment and comparable to the *Tc-cpe* knockdown. It is, however, not clear whether this is due to a knockdown of additional isoforms (as both confirmed isoforms should have been targeted by fragment 1 only as well (figure S1, B)) or to a stronger effect through targeting longer parts of the gene sequence.

**Figure 5:**
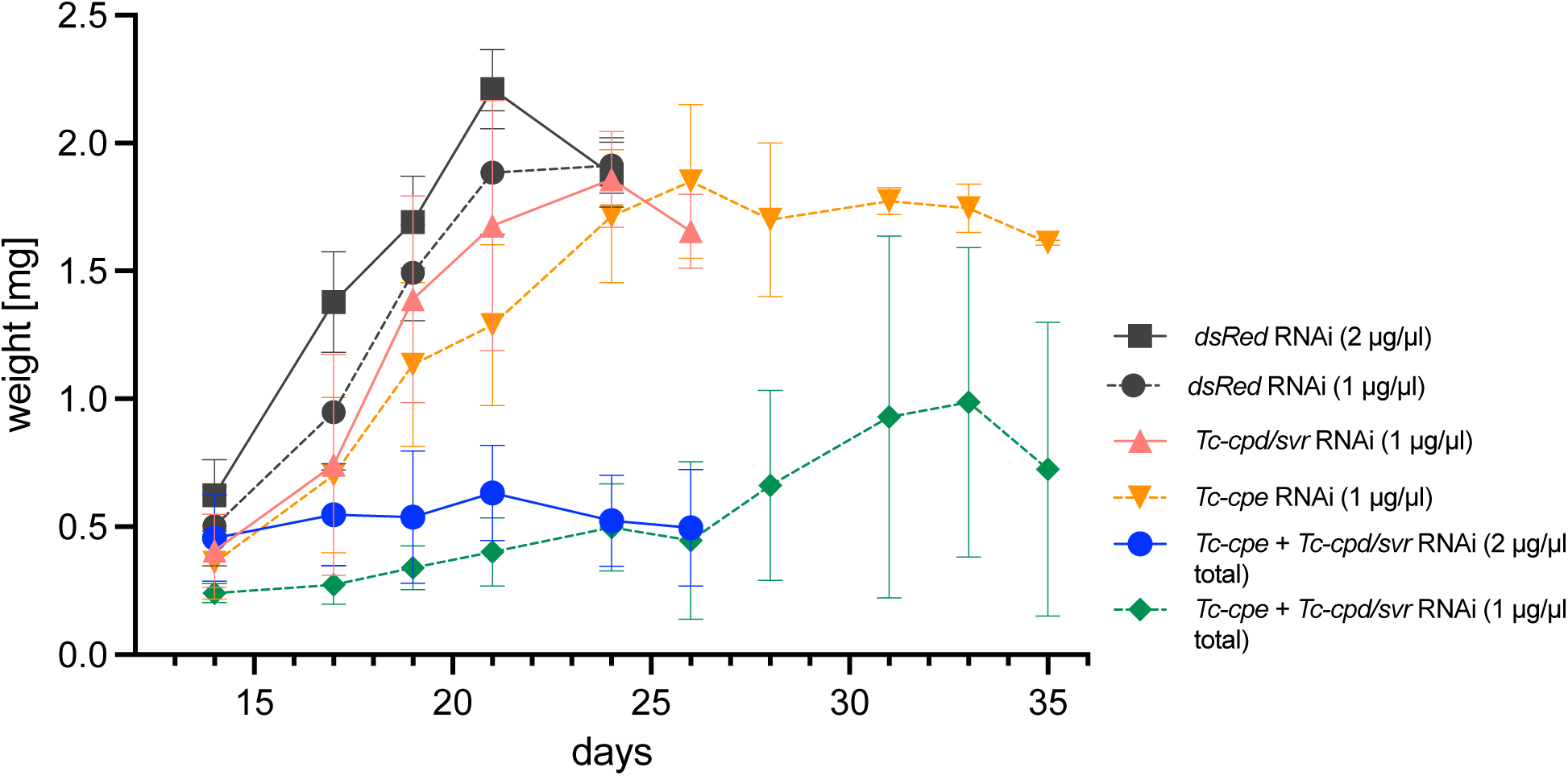
Larval growth following larval RNAi to *Tc-cpd/svr* and *Tc-cpe.* Larva were injected with dsRNA 12 days after egg lay (4^th^ larval stage) and weight was recorded from day 14 onward. DsRNA targeting this non-endogenous RNA served as a control. Weight gain curves following dsRed RNAi did not deviate from previously recorded curves on wild type samples (Fritzsche and Hunnekuhl 2021). See table 1 for sample sizes and mortality rates. Error bars indicate standard deviations. Growth curves of individual larvae can be found in figure S4.

We found a much more severe effect on growth in double knockdowns targeting *Tc-cpd/svr* and *Tc-cpe* simultaneously, conducted at 1 µg/µl and 2 µg/µl. In both sets there was almost no effective growth until day 26 of development, and all larvae injected at 2 µg/µl died from ecdysis defects by day 28. Only two of the larvae injected at the lower concentration started to gain weight again and eventually pupated (figure 5, and see figure S4 for individual growth curves). This was probably enabled by a fading out of the RNAi-effect. In summary, *Tc-cpe* and *Tc-cpd/svr* single knockdowns have only mild effects on larval growth and viability with the *Tc-cpe* phenotype achieved by a single fragment being slightly more severe. Only a knockdown of both genes in parallel causes full larval lethality through failed ecdysis.

### 4 Pupal RNAi reveals reduced viability of *Tc-cpe* RNAi treated adults and their offspring

Next, we injected *Tc-cpd/svr* and *Tc-cpe* dsRNA into female pupae to test for gene function in adult viability, egg production and embryonic development of the offspring. First, we scored survival rates of the beetles that were injected as pupae and found that *Tc-cpd/svr* knockdown does not affect viability in comparison to control groups (see figure 6A). By contrast, knockdown of *Tc-cpe* led to a higher death rate with only 56 % of animals surviving the three-week experimental time course. A repeat experiment with 50 injected pupae per background could reproduce this finding; survival of IB control was 70 %, of dsRed control and *Tc-cpd/svr* RNAi was 68 % and of the *Tc*-*cpe* dsRNA 52 % survived over three weeks.

**Figure 6:**
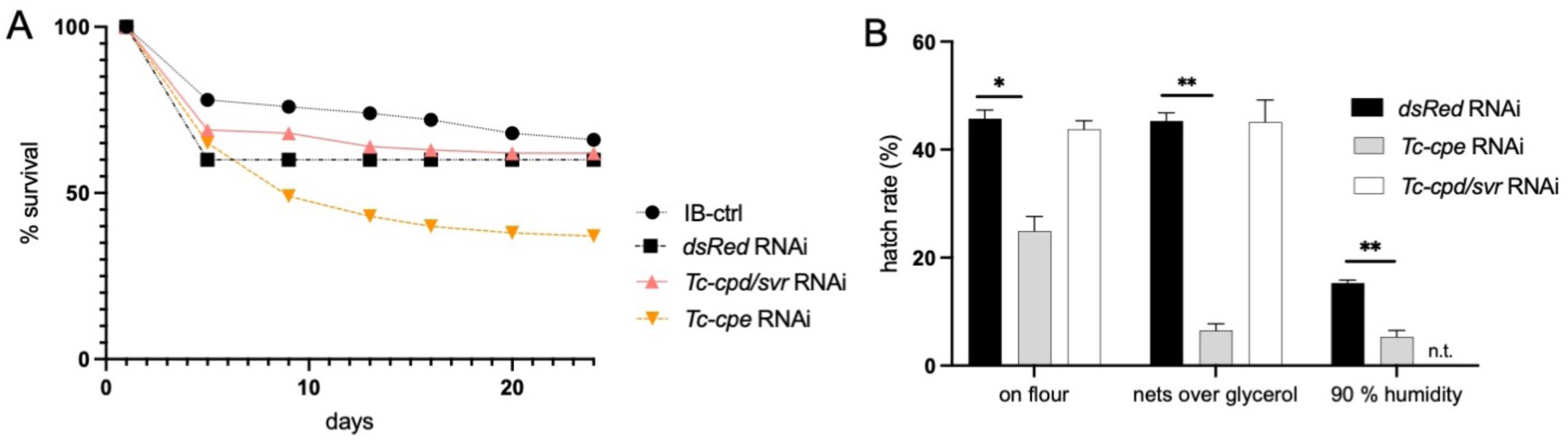
Adult survival of *Tc-cpd/svr* and *Tc-cpe*-RNAi treated female pupae (A) and hatch rates of embryos raised under different conditions (B). **A** 50 pupae of each control (IB-ctrl and dsRed RNAi) and 100 pupae of the two dsRNAs (*Tc-cpe* and *Tc-cpd/svr*) were injected. Pupae were injected at the mid pupal stage and were expected to have completed adult eclosion 5 days after injection. Deaths before day 5 were considered technical mortality and were not included in the counts (rates ranged between 20 % between and 40 % with the dsRed-control having the highest mortality rate suggesting that there is no effect *Tc-cpe* or *Tc-cpd/svr* on adult eclosion). **B** Average hatch rate/flour based on n=573 (dsRed control), n=328 (*Tc-cpd/svr* RNAi) and n=177 (*Tc-cpe* RNAi) across 3 experiments. Hatch rate/over glycerol based on n=680 (dsRed control), n=1023 (*Tc-cpd/svr* RNAi) and n=825 (*Tc-cpe* RNAi) across 2 experiments. Average hatch rate/90 % humidity based on n=424 (dsRed control) and n=369 (*Tc-cpd/svr* RNAi) across 3 experiments, not tested (n.t.) for *Tc-cpd/svr* RNAi. Error bars indicate standard deviations of hatch rates across the independent experiments, tranversal lines represent results of paired t-tests (2-tailed) with *p<0.05 and **p<0.01. No significant change (p>0.05) were found between the dsRed control group and *Tc-cpd/svr* RNAi treated specimens.

The fertility rate (defined as the number of eggs laid per female) of *Tc-cpe* RNAi treated females were reduced but only during the first 10 days after pupal injections (app. 5 days after adult eclosion). On day 9 post injection we counted in batched egglays 6.5 egg per female (n=49) in 24 h following *Tc-cpe*-knockdown, compared to 11.8 eggs per *Tc-cpd/svr*-RNAi female (n=68) and 13.3 eggs per dsRed control injected female (n=30). However, we did not observe reduced fertility of *Tc-cpe* RNAi beetles 10 days post injection and later, suggesting that the initial lower fertility observed on day 9 post injection may be an unspecific effect due to the poor condition and high mortality rate of these beetles during that period (figure 6 A). To investigate possible embryonic or early larval phenotypes by maternal transmission of the RNAi-effect to the offspring (Bucher et al. 2002), we raised eggs on 300 µm mesh width nets over Voltalef oil. This setup allows for the separation of healthy larvae which cross the net and are collected in oil from those that are affected by phenotypic defects to larval morphology. In this setup, *Tc-cpe*-RNAi larvae showed a strongly reduced hatch rate (figure 6 B). Interestingly, when attempting cuticle preparations on the sample retained on the nets, there were almost no larvae or empty eggs recovered following the lactic acid clearing protocol (Posnien et al. 2009), suggesting that eggs from *Tc-cpe*-RNAi treated mothers have a reduced resistance to chemical treatment. We were also not able to fix these embryos with our standard procedure which includes chemical treatment for dechorionization and de-vitellination with Klorix, methanol and heptane (Schinko et al. 2009). We then tested survival of the *Tc-cpe-RNAi* eggs under more natural conditions by raising them on flour. While the hatch rate was still reduced to about 50 % of what was observed for *dsRed*-control eggs and *Tc-cpd/svr*-RNAi eggs, this effect was less pronounced than in the setup over oil (figure 6 B). To test if these eggs are less resistant to increased humidity we raised *Tc-cpe*-RNAi eggs and a *dsRed-*RNAi control group in a chamber with 90 % humidity, which can normally be tolerated by the eggs (Howe 1956). While hatch rates of both samples were lower, this reduction was much stronger in the *Tc-cpe*-RNAi eggs (figure 6 B), showing that these eggs are indeed less resistant to humid conditions. By using a ubiquitous nuclear GFP-reporter line (GB233) that allowed for the easy identification of embryonic stages and developmental defects without fixation, we tested if development of *Tc-cpe*-RNAi eggs fails at a specific stage which would suggest an embryonic phenotype. We did not find any indication for developmental defects or an altered distribution of embyronic stages in a random sample of app. 30 embryos (table S3, figure S5). Hence, we conclude that the reduced hatch rate after *Tc-cpe* knock-down is due to reduced resistance to environmental influences of the eggs, presumably through an impairment of the eggshell. We then tested for an altered permeability of the eggshell for the dyes Neutral Red and Rhodamine B but found no differences to the control group (see table S4). This may indicate a reduced integrity of the chorion while the vitellin membrane is intact and cannot be penetrated by these dyes (Lemosy and Hashimoto 2000; Rand et al. 2010). In summary, we found that *Tc-cpe*-RNAi treated females have a reduced survival rate, and laid eggs with a strongly reduced hatch rate and reduced resistance to humid conditions. This is likely not due to a developmental defect but due to reduced eggshell integrity. By contrast, we did not find any effects on survival, fertility or development in *Tc-cpd/svr*-RNAi treated specimens.

### 5 Beetle *Tc-cpe* partially rescues a loss of *svr* in *Drosophila* similar to single dCPD domain constructs

*Drosophila svr^PG33^* mutants carry a P-element *Gal4* insertion upstream of the translation start in exon 1B of the dCPD-encoding gene *silver* (*svr*), which leads to a null mutation and lethality during the embryonic or early larval stages (Bourbon et al. 2002; Sidyelyeva et al. 2006). The *svr* transcript undergoes alternative splicing, leading to various dCPD variants with variable lengths (figure 1 B). The long dCPD variants carry three CP domains, of which the first two domains are active with complementary pH optima and substrate preferences, while the third CP domain is inactive (Sidyelyeva and Fricker 2002; Sidyelyeva et al. 2006) similar to vertebrate CPDs (Fricker 2025a). The lethality of the *svr^PG33^* mutants can be rescued to a variable extent by svr^PG33^-UAS-mediated expression of transgenes encoding long or short forms of dCPD that contain either all, two or only one CP domain (Sidyelyeva et al. 2010).

Since *Drosophila*, like other dipteran insects, has lost the *cpe* gene (Pauls et al. 2019), we wondered whether tCPE from *Tribolium* is functional and able to rescue a loss of dCPD in the vinegar fly. We produced UAS-*tCPE* transgenic flies by phiC integration on the third chromosome, collected males from two resulting strains (tCPE M3 and M4, figure 7 A) and crossed them to *svr^PG33^*/FM7 females. These females are heterozygous, with the recessive lethal allele *svr^PG33^* on the first (=X) chromosome rescued by a wildtype *svr* allele located on the first chromosome balancer FM7. Next, we counted the number of hemizygous *svr^PG33^/*Y (mutant) or FM7/Y (rescued control) male F1 progeny and determined the ratio of rescued flies (figure 7 B). The outcome was then compared to results from parallel experiments using UAS-rescue constructs that contain either one (*svr* 1Bshort, *svr2-3-t2*) or two (*svr*1Bt2) active dCPD domains (figure 7 A, (Sidyelyeva et al. 2010)). The ratio of rescued flies for the different dCPD rescue constructs roughly matched the results previously reported, with the highest value for *svr2-3-t2* which carries the active CP domain 2 and the inactive CP domain 3 (figure 7 B). Importantly, UAS-mediated ectopic expression of *Tc-cpe* rescued adult flies to the same extent as the native UAS-*dCPD* constructs (figure 7 A, B). During the course of our experiments, we never observed *svr^PG33^/*Y males in the *svr^PG33^*/FM7 stock, confirming that beetle tCPE is expressed and functional in the flies. Overall, our results show that, in *Drosophila*, the single CP domain of beetle tCPE can compensate for the loss of fly dCPD to a similar extent as native dCPD constructs carrying one or two active domains. We conclude from this finding that dCPD and tCPE active domains have a similar and interchangeable enzymatic activity in *Drosophila*, indicating that CPD has taken over the function of CPE in the vinegar fly.

**Figure 7:**
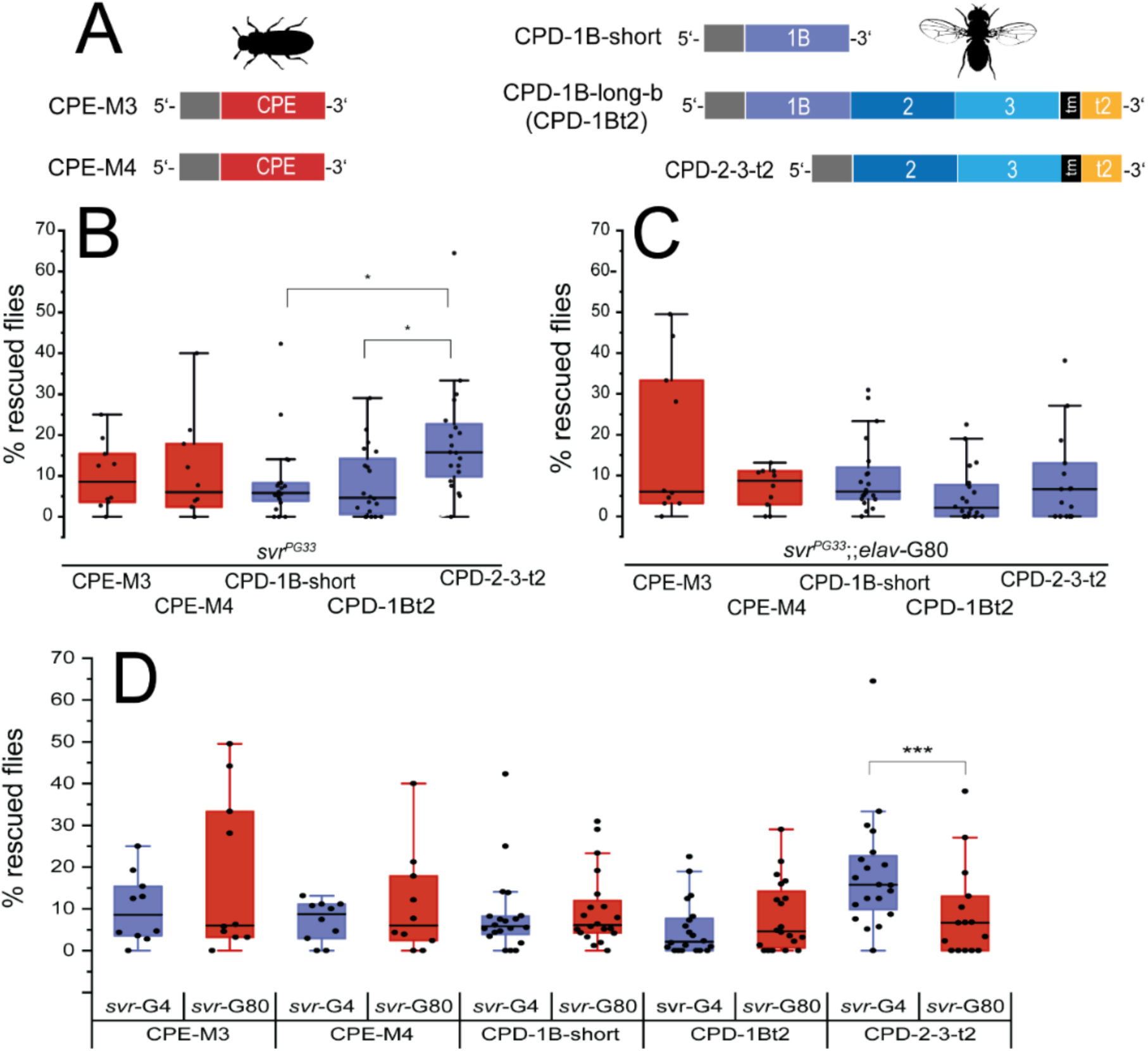
Rescue of *svr^PG33^*mutant males by expression of tCPE or specific forms of dCPD. **A** Domain structure of the used rescue constructs based on tCPE (left) and three different dCPD variants. **B** Gal4-UAS-mediated body-wide expression of tCPE (red) or specific forms of dCPD (blue) in *svr*-expressing cells, using *svr^PG33^* as Gal4-driver line. The rescue efficiency of the tested specific forms of dCPD was best for the dCPD form svr2-3-t2 (p<0.05), containing the second active CP domain and the third inactive CP domain, compared to the other tested forms (svr1B short (only first active CP domain) and svr1B-2-t2 (first and second active CP domain)). Rescue obtained with tCPE was statistically similar to any of the tested dCPD variants. **C** Gal4-UAS-mediated non-neural/non-enteroendocrine expression of tCPE (red) or specific forms of dCPD (blue) in *svr*-expressing cells, using *svr^PG33^;;elav::Gal80* (= *svr*-G80) as Gal4-driver line. Both tCPE and dCPD lines rescued to the same amount. **D** Rescue efficiency between body-wide (*svr^PG33^*, in blue) and peripheral (*svr^PG33^;;elav::Gal80* = *svr*-G80, in red) expression of tCPE or dCPD forms in *svr*-expressing cells was similar, with exception of svr2-3-t2 for which the rescue efficiency was significantly higher (p<0.001) when expressed body-wide including the nervous systems and enteroendocrine cells. Data are the same as in B and C. Pictogrammes from www.phylopic.org.

### 6 Restricted expression of beetle tCPE and fly single dCPD domain constructs is sufficient to partially rescue the loss of dCPD in *Drosophila*

Our results from *Tribolium* above showed that tCPE is mostly expressed in the nervous system (figure 4), while tCPD is believed to be more uniformly distributed in both the nervous system as well as other peripheral tissues similar to *Drosophila svr* (figure S6, S7 and (Pauls et al. 2019)). We therefore asked whether a spatially restricted ectopic expression of tCPE in peripheral tissues is sufficient to rescue the lethal phenotype of svr^PG33^ mutants. We combined *svr^PG33^* carrying a Gal4-insert in exon 1B of *svr* with the Gal4 inhibitor *elav-Gal80*. The gene *elav* is commonly considered a neuron-specific marker, even though it is expressed in enteroendocrine cells (Chen et al. 2016), and larval wing discs, fat body and trachea (Casas-Tintó et al. 2017; Winant et al. 2024). In our preparations, *elav-Gal80* largely but not completely suppressed *svr^PG33^*-driven GFP expression in the nervous system with considerable variability between individual preparations (figure S6). Peripheral expression appeared to be slightly weaker but was not removed, at least for the fat body, midgut enterocytes and epitracheal endocrine cells which we examined closer (figure S7). Next, we performed a viability assay using the different UAS-tCPE and UAS-dCPD lines as before, but driven by *svr^PG33^ elavGal80*. Again, with either tCPE and dCPD constructs, the lethality of the *svr^PG33^* allele was partially rescued to a similar extent (figure 7C). Surprisingly, the rate of rescued flies was the same for flies with whole body- or *elavGal80*-restricted expression of tCPE M3 and M4, as well as svr1Bshort, and svr1Bt2 (figure 7D), suggesting that CPD expression in many neurons is dispensable for *Drosophila* viability. Nevertheless, whole-body expression of svr2-3-t2 lead to a significantly higher rescue compared to *elavGal80-*restricted expression (Mann-Whitney, p<0.001 figure 7D). This finding suggests that the second CP domain is more effective in the CNS than in the periphery, since the svr2-3-t2 construct contains the second but lacks the first CP domain (figure 7A). In this context, it is exciting to note that we found *Tc-cpe* (carrying a CP domain homologous to the second CP domain of CPD) to be expressed exclusively in the CNS in *Tribolium*.

## Discussion

Gene loss propensity can be shaped by various evolutionary and genomic factors that influence whether a gene is retained or deleted over time. A key factor is functional redundancy arising from related genes that can compensate for the loss. For genes with redundant molecular function the propensity for gene loss is in general negatively correlated with functional importance and broad expression (Krylov et al. 2003; Makino et al. 2009; Qian et al. 2010). In this study, we showed that the genes for CPE and CPD are structurally different paralogues which can partially substitute each other in a beetle and a fly, and investigated the functional importance and distribution of *Tc-cpe* and *Tc-cpd/svr* in the beetle *Tribolium* with the aim to understand the reasons underlying the patchy gene loss of *cpe* but not *cpd/svr* in insects (Pauls et al. 2019).

Our work in *Tribolium* constitutes the first functional analysis of neuropeptide-processing carboxypeptidases in an insect that has both *cpe* and *cpd/svr*. The increased frequency of an ecdysis phenotype in *Tribolium* following *Tc-cpd/svr* and *Tc-cpe* double knockdown, which is a typical result of neuropeptide deficiencies (Rayburn et al. 2003; Rayburn et al. 2009), strongly suggests a role of both tCPE and tCPD in neuropeptide processing. Interestingly we found that *Tc-cpe* function is only partially redundant with the function of the more conserved paralogue *Tc-cpd/svr*. The strongly reduced survival of eggs, potentially through decreased eggshell integrity, that we have observed after *Tc-cpe* RNAi, was not fully buffered by tCPD function. We found that *Tc-cpe* expression first starts with the development of the nervous system, making it unlikely that a maternally transmitted dsRNA active in the embryo (Horn et al. 2022) is responsible for the increased embryonic lethality. However, the chorion, which is the outer eggshell and a barrier towards environmental influences, is secreted by the maternal follicle cells towards the end of oogenesis (Wu et al. 2008; Donoughe 2022) and hence the potential effect on this layer of the eggshell would be directly caused by gene knockdown in the mothers. While neuropeptides that could require tCPE for their processing have to our knowledge not directly been implicated in choriogenesis, egg formation is under the control of juvenile hormone and ecdysone, and different neuropeptides are known to interact with these hormonal systems (Parthasarathy, Sheng, et al. 2010; Parthasarathy, Sun, et al. 2010; Roy et al. 2018). Alternatively, tCPE could have additional functions in protein processing that are not directly connected to the neuropeptidergic pathway. The functional importance of invertebrate CPE (EGL-21) in the processing of neuropeptides has previously been demonstrated in *C. elegans*, with a deficiency of EGL-21 affecting locomotion, egg laying and metabolism (Husson, Janssen, et al. 2007).

Similar to the situation in *Tribolium*, CPE and CPD proteins are not fully mutually redundant in mice, as *Cpe^fat/fat^* mutants show defects in insulin production and a severe obese phenotype (Naggert et al. 1995) as well as strongly reduced levels of a range of different neuropeptides (Fricker et al. 1996; Che et al. 2005). These findings suggest that a considerable functional diversification of the two genes that share very similar functional domains have led to a reduced degree of redundancy in the beetle and mouse.

This poses the question what shapes the difference in gene loss propensity between insect *cpd/svr* and *cpe*. Notably, whereas genes coding for multidomain CPD-like carboxypeptidases are found across metazoans and seem to be ancient, *Cpe*-genes are restricted to bilaterians and hence appear to be evolutionarily younger (Hammond et al. 2019). Our domain-specific protein sequence analysis now showed that bilaterian *Cpe* sequences are derived from *Cpd* carboxypeptidase domain 2, strongly suggesting that the *Cpe* gene has evolved through a partial *Cpd* gene duplication in the lineage leading to bilaterians (figure 8). Moreover, unlike the ubiquitously expressed *Tc-cpd/svr* we found that *Tc-cpe* is specifically expressed in the nervous system, similar to the strong enrichment of *Cpe* in the vertebrate nervous system (Lynch et al. 1990; Fricker 2004; Ji et al. 2017). Hence, intriguingly, the evolution of an additional peptide-processing carboxypeptidase with tissue-specificity to the nervous system, as observed in these two major bilaterian groups, coincides with the origin of centralised ganglia (brains) in Bilateria. Moreover, in mammalian neurons and cell lines, CPE is enriched in “dense-core” peptidergic vesicles within the regulated secretory pathway (Docherty and Hutton 1983; Hook and Loh 1984), while CPD is enriched in the trans-Golgi network and is involved in the processing of various peptides and proteins in both the regulated and constitutive secretory pathway (Varlamov and Fricker 1998; Varlamov, Wu, et al. 1999; Varlamov, Eng, et al. 1999). Therefore, we speculate that *Cpe* has undergone subfunctionalisation and has acquired a specificity for neuropeptide/peptide hormone processing in neuronal and neuroendocrine tissues in Bilateria. The broader tissue expression as well as the broader “ancient” function in protein processing of CPD thus seems to be an underlying reason why *cpe* but not *cpd/svr* was lost in Hymenoptera and Diptera (Pauls et al. 2019). We further showed that *Tc-Cpe* can substitute the full loss of *Drosophila svr* to the same degree as single domain *svr-*constructs, indicating a high degree of molecular functional conservation between the paralogues and individual domains. However, for achieving highest rescue efficiencies a three domain *svr*-construct is necessary, showing that the full dCPD protein has the highest functionality (Sidyelyeva et al. 2010). The two active CPD domains (domain 1-2) differ in their pH optima as well as preferred cleavage site (R vs. R and L) (Fricker 2025a). CPD is therefore able to process a broader range of peptide and protein substrates than CPE, and may work efficiently in different cellular compartments with different pH (Fricker 2025a) while CPE at least in vertebrates has a pH optimum around 5.5. Moreover, at least in *Drosophila* but perhaps also other insects, peptidergic secretory vesicles have a nearly neutral pH in contrast to vertebrates (Sturman et al. 2006). The functional versatility of CPD thus might be a further reason why *cpe* but not *cpd* was lost in Hymenoptera and Diptera (Pauls et al. 2019).

**Figure 8:**
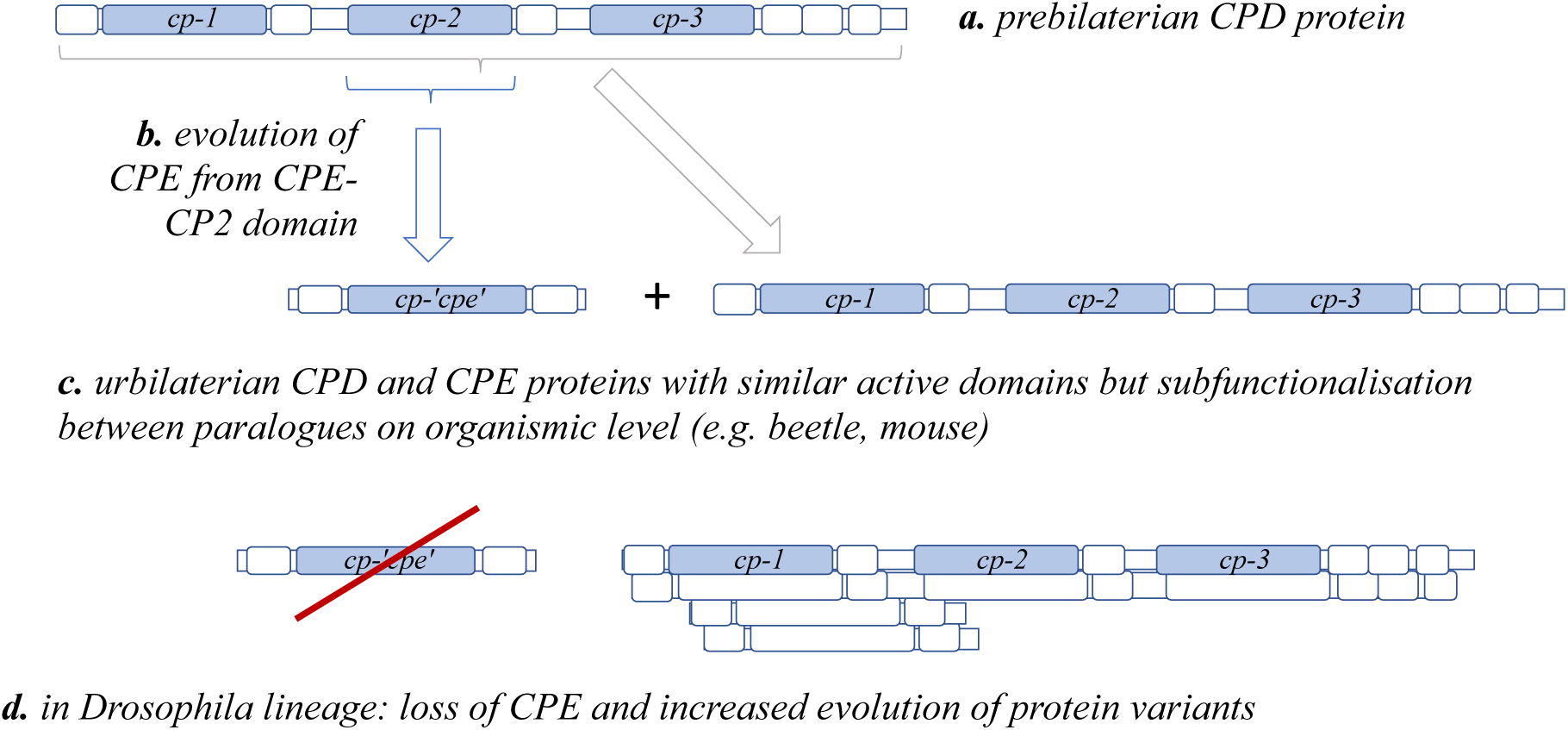
Evolution of bilaterian *Cpe* genes originating from a partial duplication of *Cpd* that included the CPD-CP2 domain (a.-c.), and loss of CPE in *Drosophila* along with an increased evolution of splice variants (d.) (also see figure S2).

Additional experimental evidence for a subfunctionalisation of CPE and larger functional versatility of CPD comes from our rescue experiments where we differentiated between a whole body (pan-) rescue and a biased peripheral rescue that excluded many neurons. Strikingly, the rescue construct that included CP-domain 2 as the only active domain showed a significantly better efficiency in the pan-rescue that included the nervous system. This is interesting with respect to our finding that CPD domain 2 is most closely related to nervous system specific CPE. Therefore, the increased efficiency of CPD domain 2 in the pan-rescue may indicate some specificity of this domain to the nervous system that could either have predated the evolution of *Cpe*, or alternatively, may have evolved along with the loss of *cpe* in dipterans. Surprisingly, we further found that despite the high abundance of neuropeptides in the nervous system, the peripheral rescue in developmental ecdysis effects was equally efficient for most rescue constructs. This might be explained by our finding that the *svr^PG33^* reporter is expressed in the epitracheal cells that produce ecdysis-triggering hormone, a key hormonal signal that is required to initiate ecdysis (Park et al. 1999).

A further reason that allowed for the loss of *cpe* in the dipteran lineage might come from the increased capability of mRNA isoform formation leading to different protein variants as observed in *Drosophila*, but to a lesser degree in *Tribolium* (see figure 1, S2). For example, the C-terminal cytosolic tail of CPD leads to enrichment of the enzyme in the trans-Golgi network (Varlamov and Fricker 1998; Varlamov, Wu, et al. 1999; Varlamov, Eng, et al. 1999). Hence, the splice variants lacking this C-terminal tail like *Dm*-CPD-1B-short might enrich the localisation to secretory vesicles, similar to CPE.

As the *cpe* gene is not only lost in dipterans, but also in hymenopterans, we checked available transcriptome data of some hymenopterans (*Nasonia vitripennis* and *Apis mellifera*, queried 07/2026 on https://www.ncbi.nlm.nih.gov/gdv) but could not find indications for multiple *cpd/svr*-isoforms. As the prediction of gene models may not reveal all possible variants, this will need additional verification. However, our current interpretation is that increased alternative splicing has not contributed to the independent loss of *cpe* in hymenopterans.

In conclusion, our results provide experimental and phylogenetic evidence that the patchy loss of *cpe* during insect evolution was possible by the structural versatility and broader intracellular and tissue expression pattern of CPD. This allowed CPD to compensate for the functionally important CPE action in the CNS.

## Materials and Methods

### Gene and protein nomenclature

We named the *Tribolium* genes *Tc-cpe* and *Tc-cpd/svr* to reflect their relationship to both vertebrate *Cpd* and *Drosophila silver (svr).* We refer to the corresponding proteins by tCPE and tCPD, in analogy to dCPD (the protein encoded by *Drosophila svr*). When referring more generally to insect *cpe* and *cpd/svr* genes we also use conventional lowercase writing. By contrast, the vertebrate genes are capitalized (*Cpe* and *Cpd*) and we also use this convention when referring to these genes more generally across animals. Protein names are always in full capitals (CPE and CPD).

### Gene and protein sequence analyses and phylogeny

*Tribolium castaneum* and *Drosophila melanogaster* gene and protein sequences were obtained through ibeetle base (Dönitz et al. 2015) and flybase (2025/4 /(Larkin et al. 2021)) respectively. The *Tribolium cpd/svr* gene models were evaluated by PCR from cDNA using specific primers which are mapped in figure S1 C and listed in table S1.

Protein features including conserved domains, PROSITE motifs, and signal peptides were obtained from UniProtKB entries through Geneious Prime 2026.0.2 (Kearse et al. 2012; The UniProt Consortium 2019). For the phylogenetic analysis, protein sequences were aligned using MUSCLE (Edgar 2004). PhyML with an LG substitution model (Le and Gascuel 2008; Guindon et al. 2010) was then used for protein tree reconstruction.

### Beetle lines

All beetles used for injections and as controls as well as for mRNA extractions and stainings were of the San Bernadino (SB) wild-type strain. To screen for possible developmental defects afer *Tc-cpe* knockdown, we used the transgenic line GB233 (nuclear eGFP-H2A fusion driven by ubiquitous RPS.3 promoter).

### *Tribolium* gene cloning, dsRNA and *in situ* probe production

Gene fragments were amplified from cDNA. Non-overlapping fragments used for dsRNA and probe production are mapped in figure S1 C and listed in supplementary table 1. For the tCPE-rescue construct the entire coding sequence of *Tc-cpe* was amplified (see table S1 for primer sequences). Products were inserted into a pJet1.2 cloning vector. For *in situ* probe synthesis the Ambion 5X Megascript T7 kit was used according to the manufacturers protocol. DsRNA fragments were produced as described in (Posnien et al. 2009).

### *Tribolium* RNA *in situ* hybridisations

Embryo fixation of mixed stages ranging from 0-72 hours of development and mRNA in situ labelling were carried out as described (Schinko et al. 2009). RNA stainings in dissected nervous systems along with monoclonal anti-synapsin antibody (mouse 3C11, DHSB Hybridoma Bank) labelling of neuropile and DAPI staining of nuclei were performed as described (Hunnekuhl et al. 2020).

### *Tribolium castaneum* larval and pupal RNAi experiments

Larval RNAi followed by the assessment of survival, frequency of ecdysis phenotypes and larval weight at different timepoints was carried out as described in (Fritzsche and Hunnekuhl 2021). For parental RNAi female pupae were injected with dsRNA as described in (Posnien et al. 2009). They were crossed to wildtype males directly after adult eclosion. App. 7 days after injection 24 h egg-lays were set up and eggs were collected by sieving. Eggs were raised at 32°C in small, otherwise emtpy plastic vials, on 300 µm nets over glycerol or in a closed chamber in which an open beaker with water was placed as well. Humidity in this chamber was recorded and constantly reached 90 %. For cuticle analyses eggs were left to develop for 72 h and then transferred into lactic acid/ 10% ethanol on slides, topped with a cover slip and left to clear overnight at 60°C.

### *Tribolium* eggshell permeability and developmental assays

For assessment of possible developmental defects, female pupae of the GB233-nuclear GFP reporter line were injected with 1 µg/µl dsRNA targeting *Tc-cpe* or non-endogenous dsRed. Females were crossed to San Bernadino wildtype males and embryos were collected afer eclosion. Embryos were washed briefly in 1 % Klorix to remove flour and were embedded without fixation in 80 % glycerol for micrsocsopic inspection using a Zeiss LSM980 confocal microscope.

To test for a possible increased permeability of eggs collected after pupal *Tc-cpe* RNAi treatment, we used Neutral Red (Lemosy and Hashimoto 2000) and Rhodamine-B (Rand et al. 2010) assays. For both assays 70 RNAi treated females (1 µg/µl *Tc-cpe* dsRNA/ 1 µg/µl dsRed RNAi) were put on cages over filter paper to ensure flour free egg collection as described (Gautam et al. 2015). Embryos were collected the next day and washed two times in PBT. For Neutral Red staining embryos were incubated in 5 mg/ml Neutral Red (Thermofisher N3246) in PBT for 20 minutes. Embryos were then washed 2 x with PBT again.

For Rhodamine-B staining, embryos were incubated for 15 minutes in a 1 mM/PBT Rhodamine-B (Sigma-Aldrich R6626) solution and re-transferred into PBT. Embryos of both treatments were observed under a standard stereomicroscope with external light source and with a LEICA M205 FA epifluorescence stereomicroscope (Rhodamine-B assay).

### Quantitative PCR assays

Primers used in the qPCR are listed in table S1. For the *Tc-cpd/svr* and fragments are indicated in figure S1C. *Tc-cpe* gene expression assays mRNA was extracted from the specific stages using the Quick-RNA Tissue/Insect Kit including on-column DNAseI digest (Zymo Research), and subsequently reverse transcribed to cDNA using SuperScript III (Invitrogen) and oligo dt primers. Concentrations of the samples were adjusted to equal values. Quantitative PCR was then conducted as described (Fritzsche and Hunnekuhl 2021). Briefly, primer pairs with approx. 120 bp distance were tested for adequate PCR efficiency using a cDNA dilution series. The qPCR experiments were run using a C1000 thermal cycler with a CFX96 detection system (Bio-Rad) with technical triplicates for each sample. Data were analysed using the comparative CT method (Schmittgen and Livak 2008). For assessment of knockdown levels following RNAi, mRNA extracted from larvae 24 h after injection was extracted, reverse transcribed into cDNA and used in qPCR with the same primer pairs as above. A primer pair targeting the ribosomal protein RPS3 served as a standard (table S1). Statistical tests (paired t-tests and standard deviations) were applied to the fold change in 3 biological using GraphPad Prism 11.0.2 (GraphPad Software, Boston, MA, USA).

### Fly lines

The following fly lines were used: *y^1^ w* P{w^+mWhs^=GawB}svr^PG33^*/FM7h (*svr^PG33^*/FM7), *w*;UAS:svr1*B-short, *w*; UAS:svr*1B-2-3-t2 (1Bt2), *w*;UAS:svr*2-3-t2 (kind gift of Galina Sidyelyeva and Lloyd Fricker (Sidyelyeva et al. 2006; Sidyelyeva et al. 2010)), w*; P{w^+mC^=UAS-mCherry.NLS}3 (BDSC #84277) and w*; P{y^+t7.7^ w^+mC^=10XUAS-IVS-myr::GFP}attP40 (BDSC #32198). We further generated a stable *y^1^ w* P{w^+mWhs^=GawB}svr^PG33^*/FM7h;;*elavGal80* (*svrPG33 elavG80*) line combining *svr^PG33^*/FM7 and *w*;; P{w^+m*^=elav-GAL80.S*}3 (BDSC #602882).

### Generation of UAS-tCPE flies

The *tCPE* sequence was amplified by PCR from *tCPE* cDNA with a Phusion high fidelity polymerase (Thermo Fisher) and the primer set 5’-CCGCGGCCGCCAAAATGGTGTTCGCAACATTGTTG-3’ (containing a *NotI* restriction site and a Kozak optimisation sequence) and 5’CGTCTAGATTATTACAGATCTTCTTCAGAAATAAGTTTTTGTTCTTCAACTGCTATCGGTTGCCTG-3’ (containing a stop codon, Myc tag and a *XbaI* restriction site). The amplicon was restriction-cloned into the pUASTattB vector (kind gift of Konrad Basler (Bischof et al. 2007)), and the sequence verified by Sanger sequencing (supplementary Material S1). Transgenic flies were produced by BestGene Inc. (Chino Hills, CA, USA), based on phiC31 integrase-mediated transformation in y^1^ w^67c23^; P{y^+t7.7^=CaryP}attP2 (BDSC # 8622) flies.

### Fly survival assay

Females of the *svr^PG33^* or *svr^PG33^;; elav* G80 line were crossed with males of the various UAS-dCPD or UAS-tCPE constructs. Per vial, five virgin females and five males were combined and flipped once a week for a total of 3 weeks. For each fly line, 10 vials (N=10) were established, and the resulting adult males were scored for *svr^PG33^*/Y or *svr^PG3^;;elavG80*/Y or FM7/Y, based on the Bar (B) eye phenotype. The ratio of rescued flies was calculated as # (*svr^PG33^*/Y males)/(# (*svr^PG33^*/Y males) + # (*svr^PG33^*/FM7 males)).

## Statistics

The *Drosophila* rescue experiments were analysed by a Kruskal-Wallis-ANOVA with Dunn’s correction (comparison of all strains) or Mann-Whitney U test (pairwise comparison) in OriginPro, Version 2023b (OriginLab Corporation, Northampton, MA, USA). For the *Tribolium* data standard deviations and pairwise two-tailed t-tests (for fig.6 B) were computed using using GraphPad Prism 11.0.2 (GraphPad Software, Boston, MA, USA).

## Acknowledgment

We thank Claudia Hinners and Gertrud Gramlich for excellent technical assistance, Konrad Basler, Lloyd Fricker and Galina Sidyelyeva for the kind gift of plasmids and flies, Sonja Fritzsche for supporting some initial *Tribolium* experiments, and Gregor Bucher for sharing laboratory infrastructure and beetle stocks.

## Author contributions

VSH and CW conceived the study, VSH conducted the gene phylogenetic and transcript analyes, VSH and JCH performed the *Tribolium* experiments and data analysis, CW performed the *Drosophila* experiments and data analysis, VSH and CW wrote the paper, all three authors read and approved the final manuscript.

## supplementary materials

**Figure S1:**
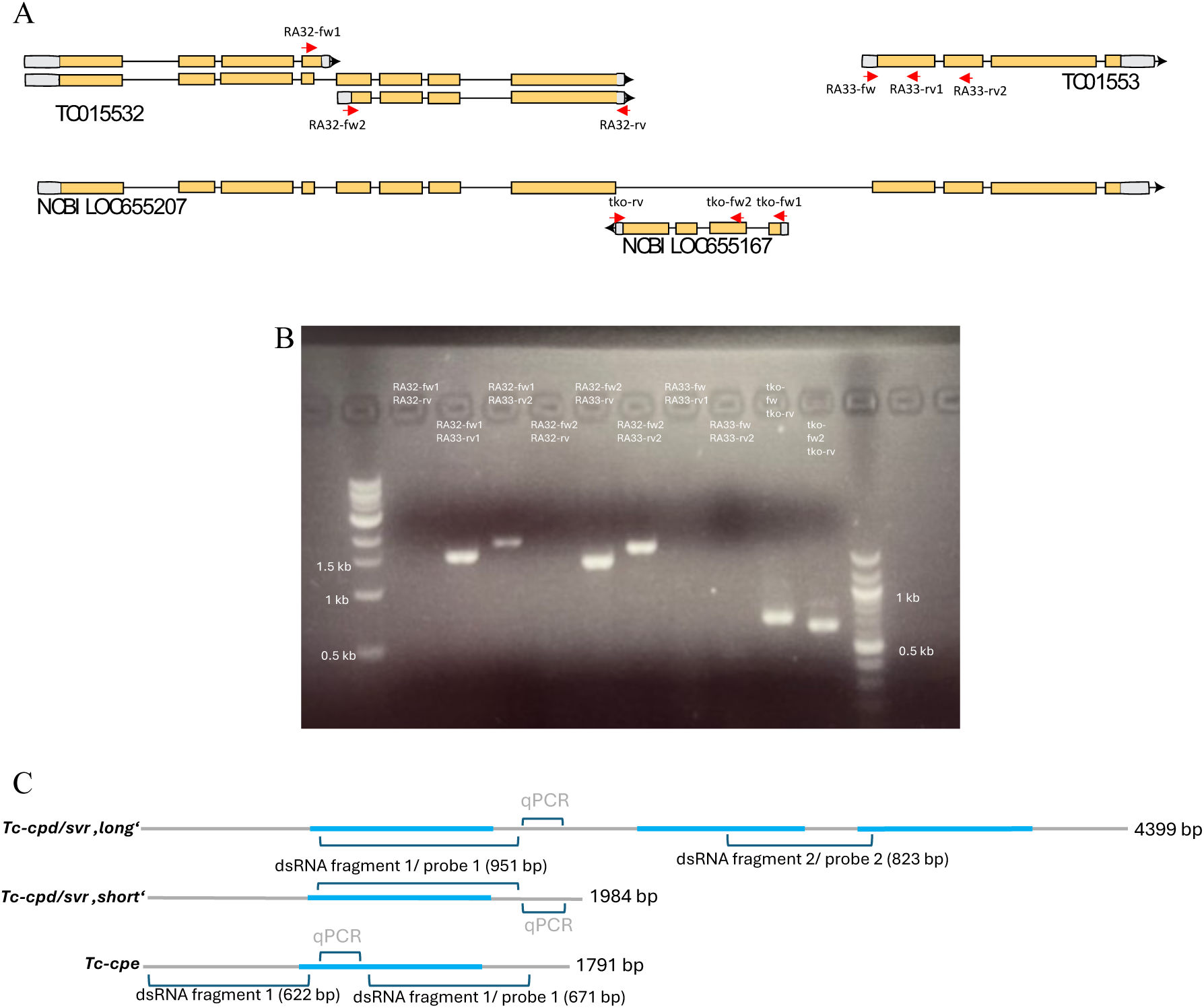
A) Gene models in the *Tribolium cpd/svr* locus (from https://ibeetle-base.uni-goettingen.de/) B) PCR validation of predicted models from (A), C) Maps of PCR-validated *Tribolium* Cpd and Cpe mRNAs including fragments used in RNAi, qPCR and in situ hybridisation experiments. Primer sites that we used to test for the putative presence of additional predicted transcripts did not produce bands (A/B), but we could retrieve fragments (B) that confirmed the long transcript spanning the annotations TC015532 and TC015533, which were wrongly interpreted as two separate genes. Nested in the large 8^th^ intron of the full *Tc-cpd/svr* gene but in reverse orientation is a *takeout-like* gene (NCBI LOC655167) and part of the 3-prime end of this gene have wrongly assigned to the partial *Tc-cpd/svr* transcripts. Brackets in C) indicate the gene fragments that were used in RNAi (dsRNA) and *in situ* hybridisation (probes) experiments and amplified for the qPCR assays (Figure 3 and S3) blue: sequence encoding peptidase functional domains.

**Figure S2:**
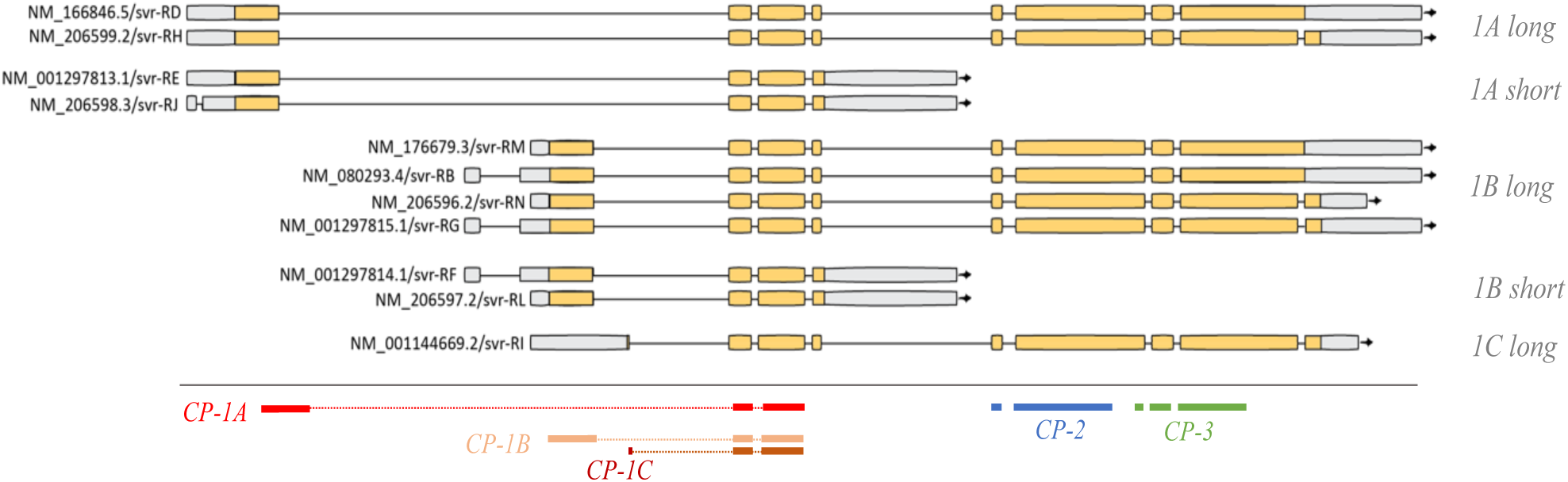
*Drosophila svr* gene models, retrieved from flybase/*D. melanogaster* r6.65 genome, models based on RNA sequence evidence. Sequence span encoding the carboxypeptidase active domains are mapped in bottom lines.

**Figure S3:**
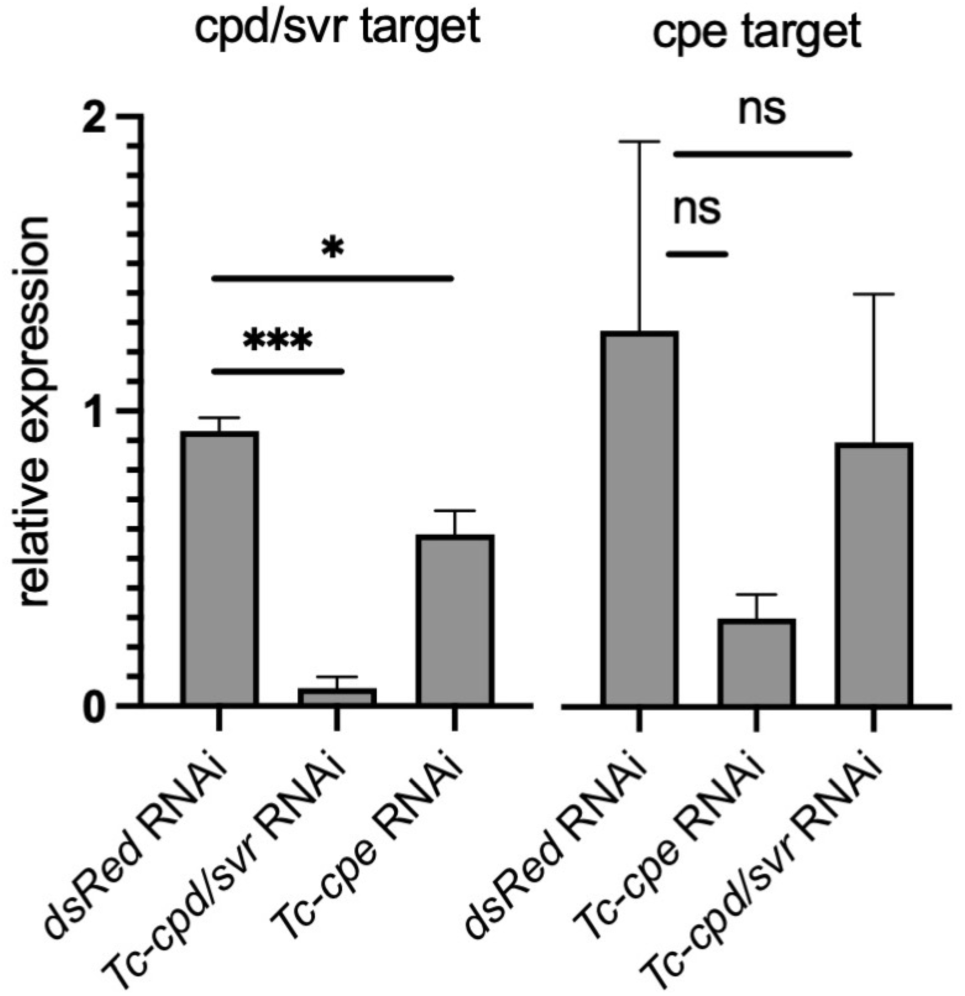
Assessment of knockdown efficiencies and test for cross-reactivity. As two independent controls RNA from dsRed-RNAi. *Tc-cpe* expression was considerably reduced to app. 30 % of the WT-expression in the *cpe*-RNAi background, although the change in relation to the dsRed control sample was just above the conventional significance threshold of 0,05 (paired two-tailed t-test P=0.059). *Tc-cpd* expression was severely reduced to app. 6 % of the WT expression and the reduction was highly significant in relation to the dsRed control sample. We did not observe large effects of the knockdown in the expression of the non-target paralogue. Expression levels were normalized to the expression of RPS.3 and expression was analysed in relation to wildtype samples (≙1) using the comparative CT method (Schmittgen and Livak 2008). Error bars indicate standard deviations with only positive values shown. Lines represent paired t-test results (2-tailed): ns (not significant p>0.05): p>0.05, *= p<0.05, ***=p<0.001.

**Figure S4:**
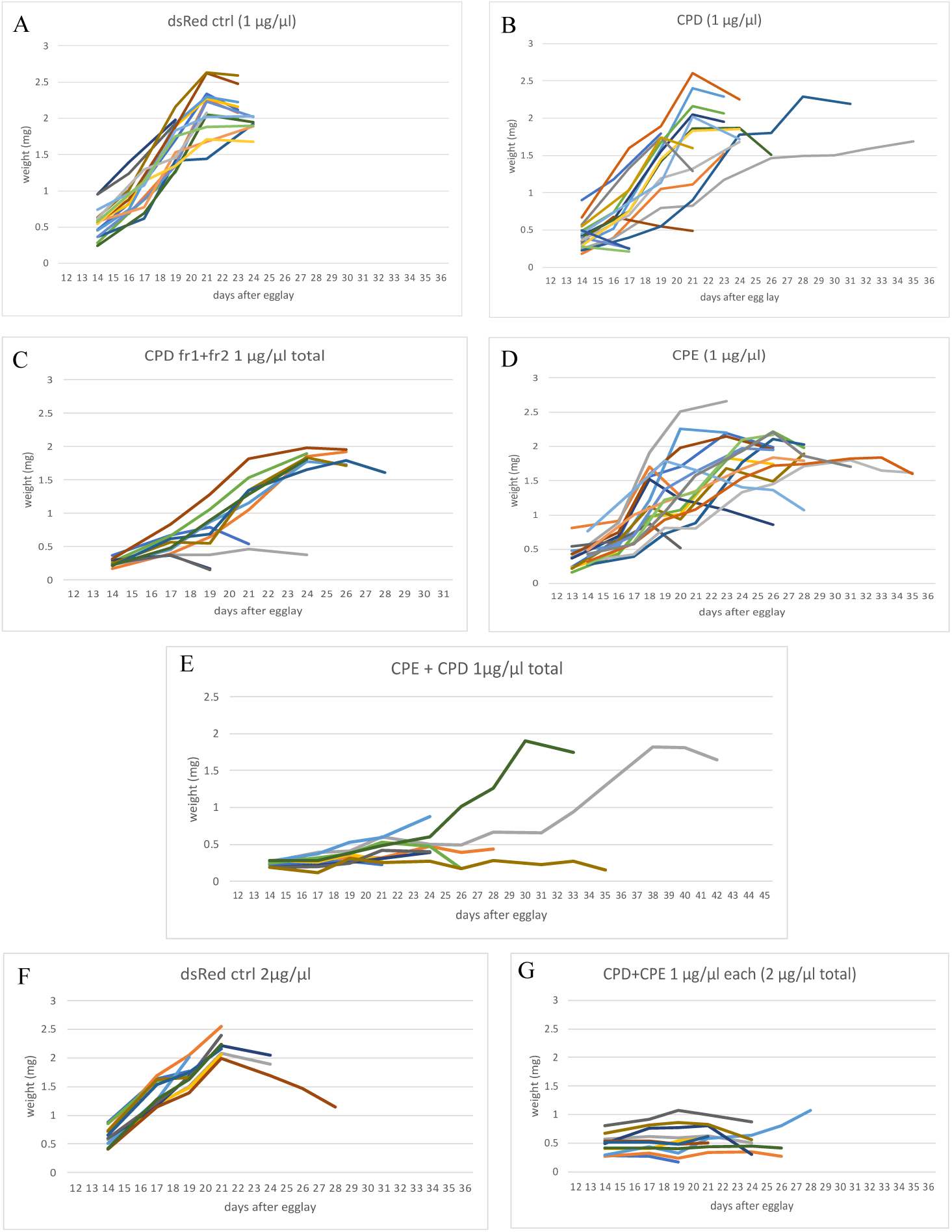
Individual larval growth curves following dsRNA injections. Data from A-D and F-G is basis for composite curves in figure 5 /main manuscript. Each coloured line represents individual larvae that was injected with the respective treatment at day 12 of development. **A** n=18, **B** n=22, **C** n=12, **D** n=22, **E** n=12, **F** n=12, **G** n=12.

**Figure S5:**
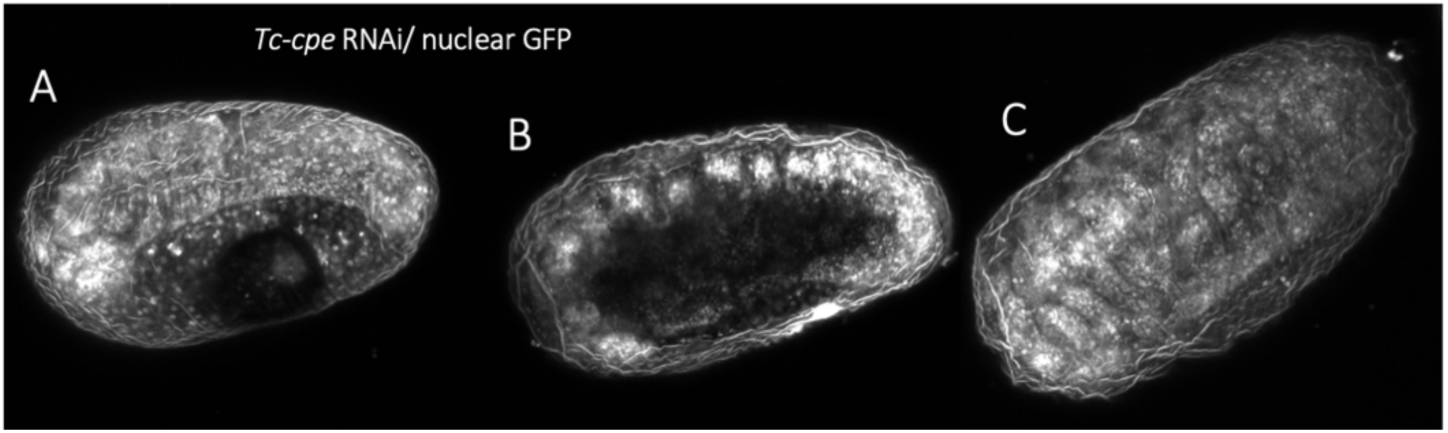
Examples for stage categories, all from the Tc-cpe RNAi background. A) early germband, B) elongated germ band, C) late embryo with developed appendages. Line: GB233 (ubiquitous nuclear GFP).

## Material S1 translated sequence of tCPE, after sequencing the final transformation vector (pUASTattB with tCPE insert)

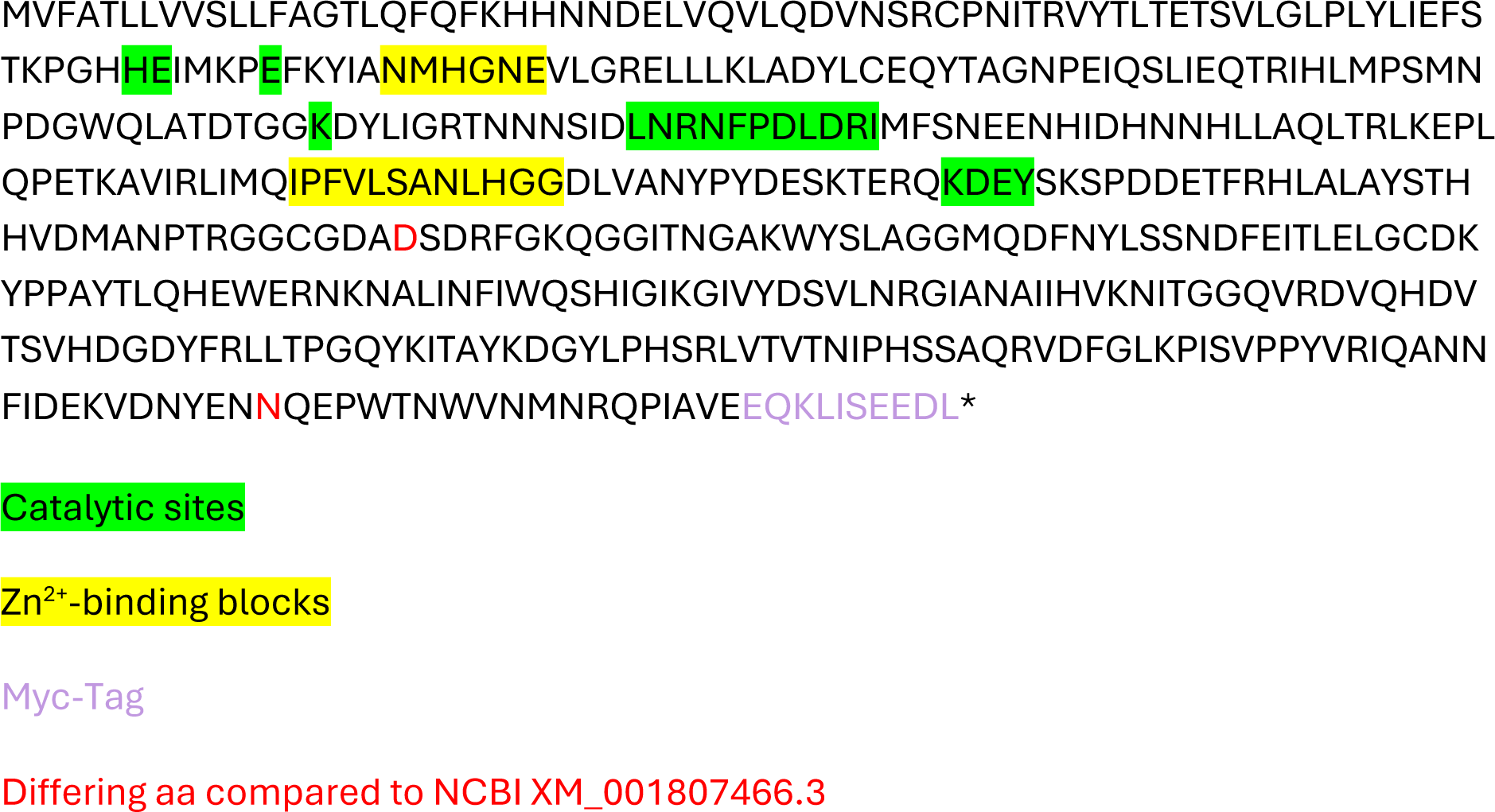

---

**Figure S6:**
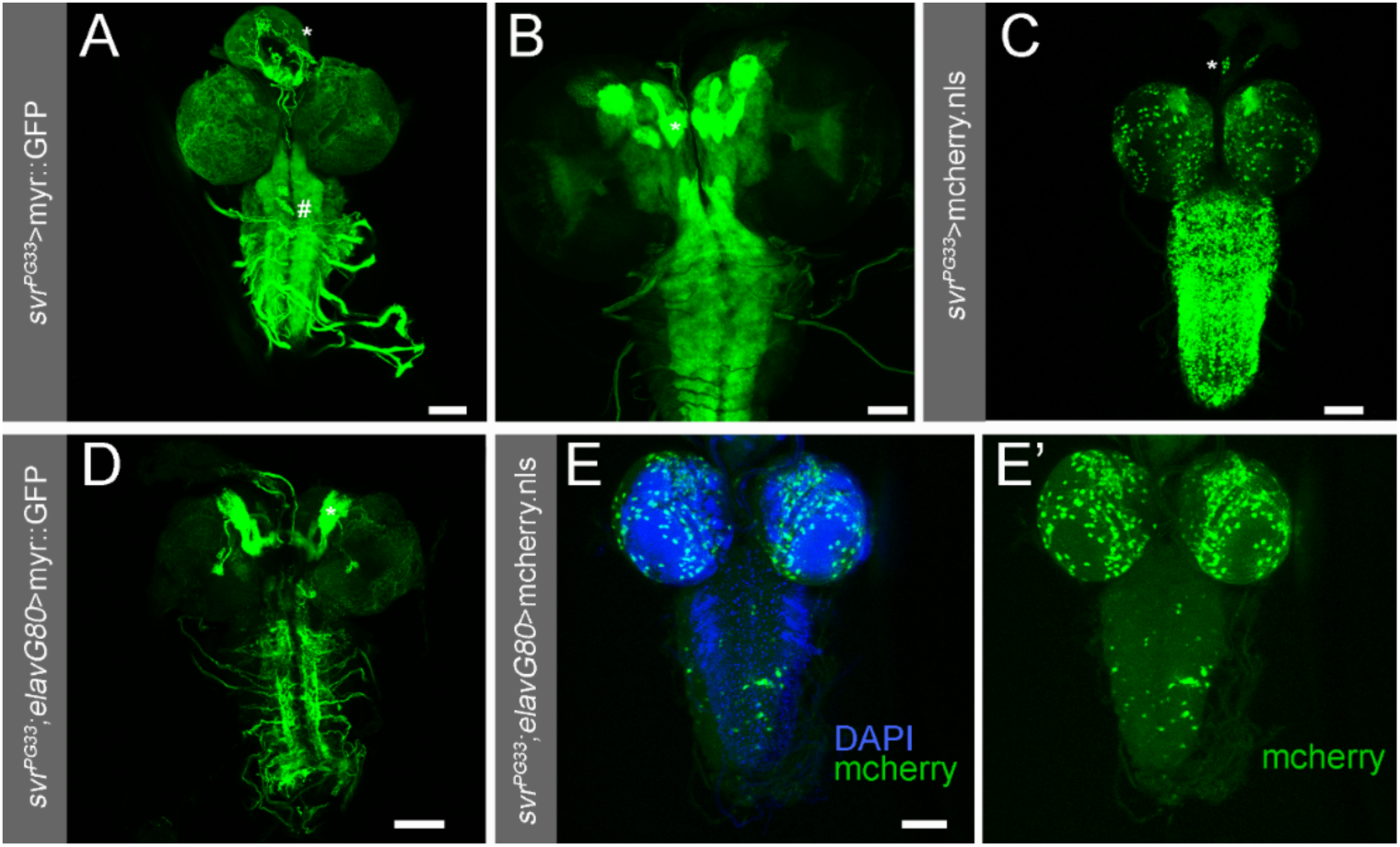
Expression pattern of *svr^PG33^* in the L3 larval nervous system of *Drosophila*. The PG^33^ Gal4-insertion is located in exon 1B of *svr* and hence is expected to reflect the native *svr* expression pattern. **A** PG^33^-mediated expression of membrane-bound *myr::GFP* labels particularly the ventral nerve cord and neurosecretory terminals in the ring gland (*) and thoracic and abdominal perisympathetic organs (#). Full confocal stack **B** Confocal stack comprising the mushroom body area from a different preparation. The mushroom body calyx and peduncle is visible in each brain hemisphere. **C** PG^33^-mediated expression of nuclear mCherry broadly labels nuclei in the ventral nerve cord, with more restricted labelling in the brain lobes particularly in the Kenyon cells of the mushroom body. Also the endocrine adipokinetic-hormone producing cells in the ring gland are labelled (*). **D** Introduction of *elavGal80* strongly but not completely reduced the expression of *PG^33^*-mediated expression of GFP in the L3 nervous system. In the brain, the mushroom bodies are still strongly labelled. **E-E’** *elavGal80* strongly reduced the *PG^33^*-mediated expression of nuclear mCherry (green) in the ventral nerve cord, and to a lesser extent in the brain. Co-labelling of all nuclei by DAPI (blue) shows that only relatively few neurons express mCherry. Scale bars = 50 µm.

**Figure S7:**
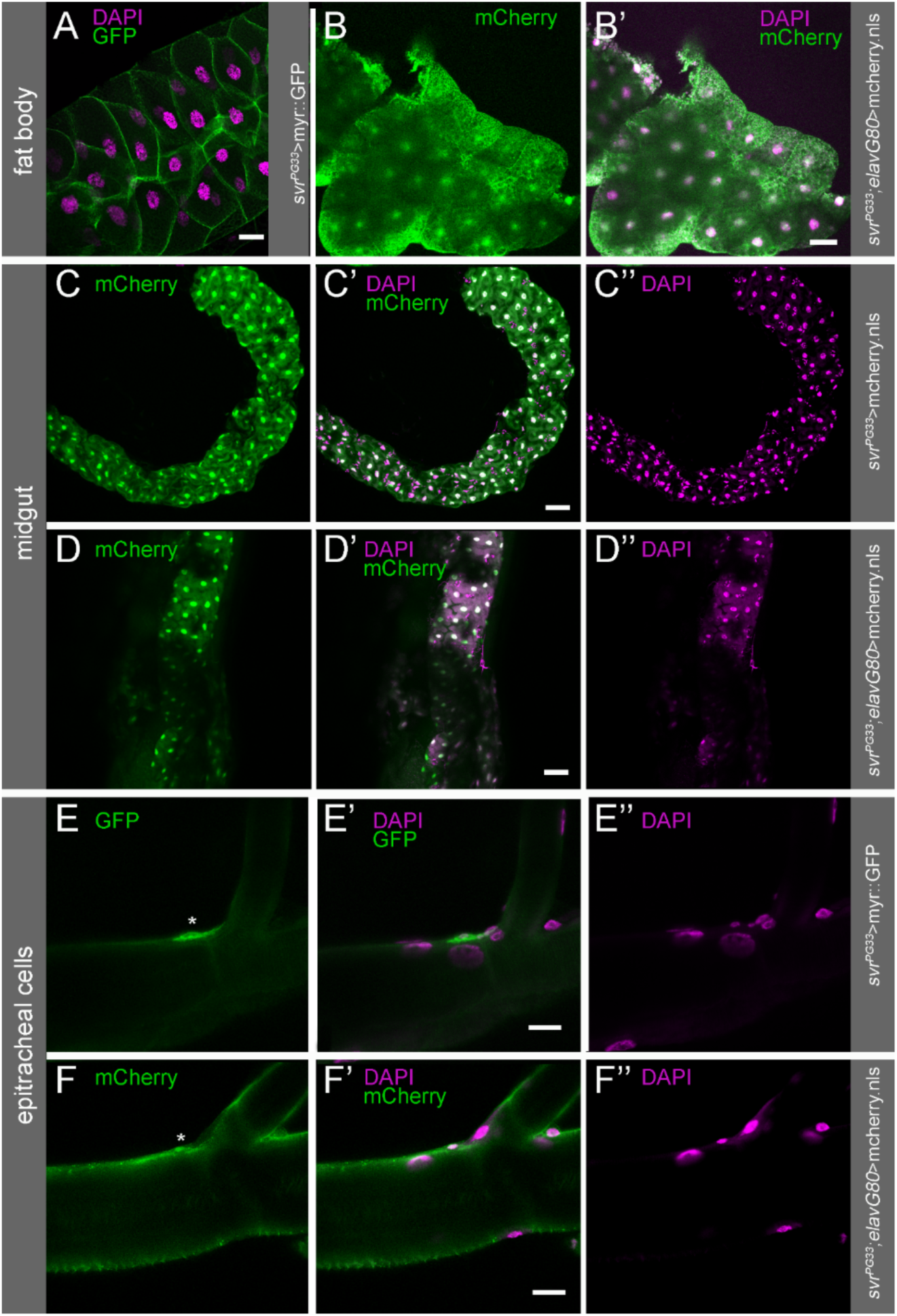
Expression pattern of *svr^PG33^* in peripheral tissues of the L3 *Drosophila*. The PG^33^ Gal4-insertion is located in exon 1B of *svr* and hence is expected to reflect the native *svr* expression pattern. **A** PG^33^-mediated expression of membrane-bound GFP (green) strongly labels fat body cells. Nuclei are stained by DAPI (magenta). **B-B’** *elavGal80* does not suppress PG^33^-mediated expression of nuclear mCherry (green) in the trophocyte nuclei stained by DAPI (B’, magenta). **C-C’’** PG^33^-mediated expression of nuclear mCherry (C-C’, green) strongly labels enterocytes and enteroendocrine cells, but not dividing stem cells. Nuclei are stained by DAPI (C’-C’’, magenta). **D-D’’** *elavGal80* does not suppress PG^33^-mediated expression of nuclear mCherry (D-D’, green) in the midgut. Nuclei stained by DAPI (D’-D’’, magenta). **E-E’’** PG^33^-mediated expression of membrane-bound GFP (E-E’, green) labels the endocrine epitracheal cells (asterisk, only one segment shown). Nuclei are stained by DAPI (C’-C’’, magenta). **F-F’’** *elavGal80* does not suppress PG^33^-mediated expression of nuclear mCherry (green) in the epitracheal cells (asterisk, only one segment shown). Nuclei stained by DAPI (F’-F’’, magenta). Scale bars = 50 µm (A-D’’) and 20 µm (E-F’’).

**Table S1:**
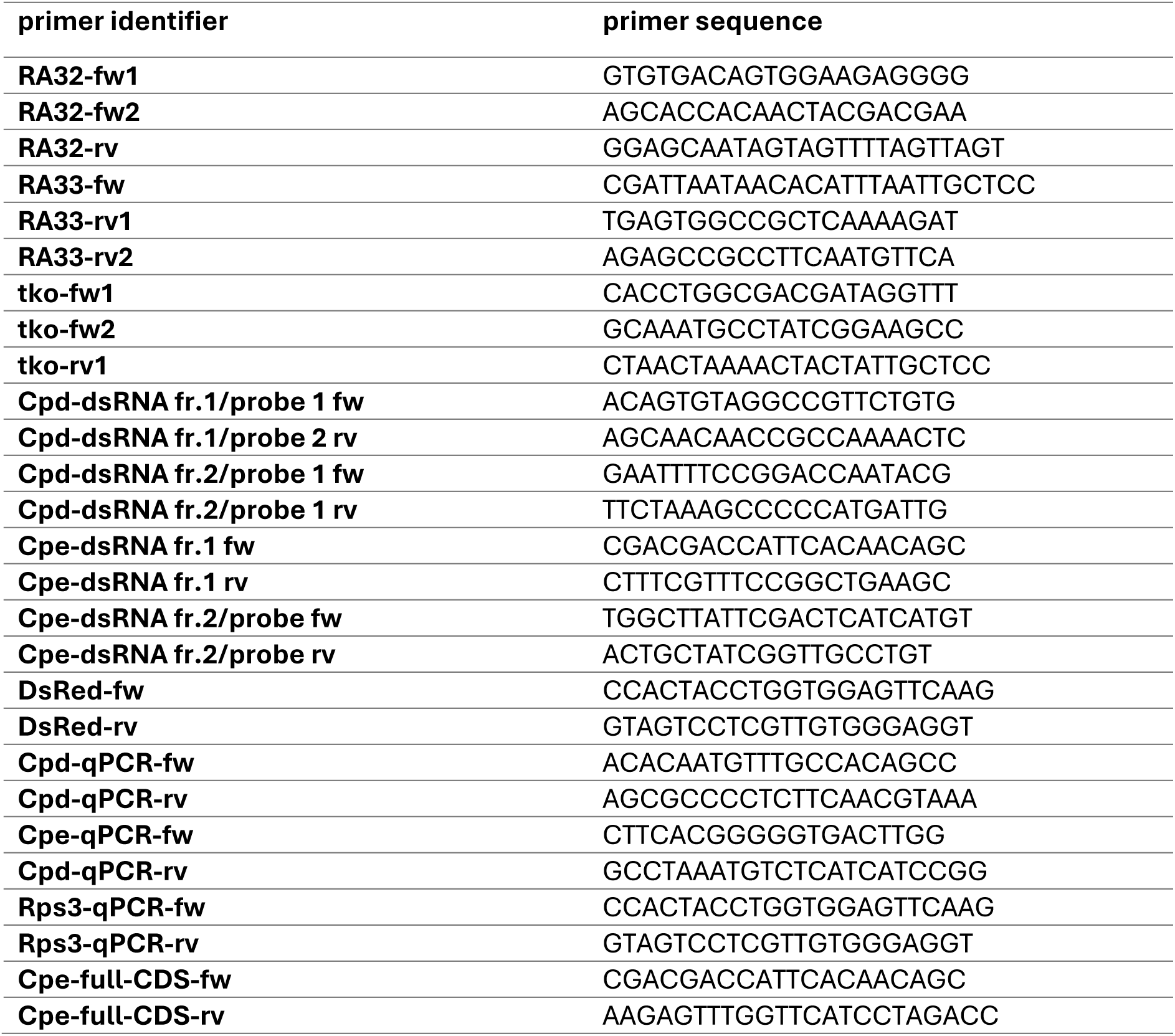
Primers used to amplify *Tribolium castaneum* gene fragments and in the qPCR assays.

**Table S2:** Classification of *Drosophila melanogaster* silver proteins based on translation of mRNA isoforms. Variants can be grouped into 5 major classes depending on the version of the first CPD domain (A/B/C) and the presence of all three active domains (long variants) or only carboxypeptidase domain 1 (short variants). Some individual variants are encoded by two mRNAs with differing UTRs. The versions CPD-1A-long-a and -b and CPD-1B-long-a and -b only differ in length and sequence of the cytosolic tail following the transmembrane domains.

| protein variant | length | NCBI transcript acc.# |
| --- | --- | --- |
| CPD-1a-long-a | 1456 AA | NM_166846.5 |
| CPD-1a-long-b | 1423 AA | NM_206599.2 |
| CPD-1a short | 452 AA | NM_001297813.1; NM_206598.3 |
| CPD-1b- long-a | 1439 AA | NM_206596.2; NM_001297815.1 |
| CPD-1b-long-b | 1406 AA | NM_176679.3; NM_080293.4 |
| CPD-1b-short | 435 AA | NM_001297814.1; NM_206597.2 |
| CPD-1c-long | 1305 AA | NM_001144669.2 |

**Table S3:** Developmental stages identified 24 – 72 h after egg lay from *Tc-cpe* RNAi treated mothers and a *dsRed*-dsRNA injected control group of the homozygous nuclear GFP-reporter line GB233. Injected females were crossed to wildtype males and randomly chosen eggs were inspected using a confocal microscope and sorted into the given stage categories they were most similar to (see figure S3). No pre-germband stages were expected, and stage distribution was comparable between both backgrounds. Not defined: stage could not be defined, possibly non-fertilized or mis-developed.

| Stage | # embryos *dsRed*-RNAi control | # embryos *Tc-cpe* RNAi |
| --- | --- | --- |
| early germband | 5 | 4 |
| elongated germband | 9 | 10 |
| late embryo with developed appendages | 15 | 13 |
| not defined | 4 | 3 |
| Total | 33 | 30 |

**Table S4:** Neutral Red and Rhodamine-B assays of egg-shell permeability.

| dye | # embryos | detection | observation after transfer into PBT |
| --- | --- | --- | --- |
| Rhodamine-B | 2 x 30 / *Tc-cpe*-RNAi | epifluorescence and stereomicroscope with external light source | No internal signal/ no fluorescent signal (equal in both backgrounds) |
| Rhodamine-B | 2 x 30 / *dsRed*-RNAi |  |  |
| Neutral Red | 2 x 30 / *dsRed*-RNAi | stereomicroscope with external light source | No internal signal, quickly fading peripheral signal, (equal in both backgrounds) |
| Neutral Red | 2 x 30 / *Tc-cpe*-RNAi |  |  |

